# Polygenic Effects on the Z Chromosome Underlie Hybrid Incompatibility in *Papilio* and *Heliconius* Butterflies

**DOI:** 10.1101/2022.10.28.514284

**Authors:** Tianzhu Xiong, Shreeharsha Tarikere, Neil Rosser, Xueyan Li, Masaya Yago, James Mallet

**Affiliations:** Department of Organismic and Evolutionary Biology, Harvard University; Cambridge, MA, 02138, USA; State Key Laboratory of Genetic Resources and Evolution, Kunming Institute of Zoology, Chinese Academy of Sciences; Kunming, Yunnan, 650223, China; The University Museum, The University of Tokyo; Bunkyo-ku, Tokyo, 113-0033, Japan

## Abstract

The fitness of animal hybrids follows two empirical rules: hybrids of the heterogametic sex are more unfit (Haldane’s Rule), and the sex chromosome is disproportionately involved in incompatibility (the large-X/Z effect). Whether these rules result from genetic mechanisms shared across taxa remains unknown, and existing explanations rarely consider female heterogametic taxa such as butterflies. Here, we investigate hybrid incompatibilities in *Papilio* and *Heliconius* butterflies, and show that defects coincide with unbalanced introgression between the Z chromosome and its genetic background. This polygenic mechanism predicts both rules because introgressed ancestry on the Z chromosome is more skewed in females, and is more variable than on all autosomes. Therefore, the explanation for both rules in butterflies shares little similarity with prevailing theories relying on dominance.

## Introduction

Speciation is a complex and stochastic process, yet it obeys empirical rules across taxa with sexual reproduction (*1*). Haldane’s Rule states that among the hybrids between different species, the heterogametic sex (the sex with XY or ZW sex chromosomes) tends to have lower fitness (*2, 3*). A second rule is the so-called large-X effect, which states that the sex chromosome is disproportionately involved in hybrid incompatibility (*1, 4*). Haldane’s Rule is entirely phenomenological, but it holds across many phylogenetically diverse organisms (*5*–*7*). Whether adherence to Haldane’s Rule emerges from a common set of genetic mechanisms is an open question (*8*). The large-X effect also appears robust, but without mapped incompatibility factors, the evidence is often indirect and hard to interpret (*9*–*13*).

Several mechanisms can explain Haldane’s Rule. First, dominance theory posits that the single X chromosome in males might expose recessive genes that are deleterious in a hybrid genetic background, thus increasing the likelihood of incompatibility (*4, 14*). Second, the evolution of sex chromosomes may be faster than autosomes, thus Haldane’s Rule can be produced via a variety of processes involving hemizygous haploid selection (*15*), sex-specific selection (*16*), and sex chromosome conflict (*17, 18*). Accelerated sex chromosome evolution also provides a natural explanation for the large-X effect. Empirical evidence for these mechanisms is based mostly on male heterogametic taxa, such as mammals and *Drosophila* (*19, 20*), but whether they are general explanations for the two rules is unknown. It has also been suggested that spermatogenesis is particularly prone to disruption, which can explain the higher incidence of male sterility in XX/XY systems (*21*), but this is not applicable to Haldane’s Rule in female heterogametic taxa.

In Lepidoptera (butterflies and moths), the female is the heterogametic sex (with Z/W sex chromosomes), and hybrid females are more prone to defects than males (*2, 22*). To date, little is known about the genomic basis of Haldane’s Rule in Lepidoptera, except for a few studies using sparse genetic markers (*23*–*25*) and a recent whole-genome Quantitative Trait Locus (QTL) study in *Heliconius* (*26*). Nonetheless, these studies demonstrate that hybrid female sterility is associated with the Z chromosome, consistent with a large-Z effect. Here, we map hybrid incompatibility between two closely related butterflies, *Papilio bianor* and *Papilio dehaanii* (Fig. 1A), in which interspecific crosses produce fertile males but completely sterile females (*27, 28*). Hybrids also develop abnormal body size, which is frequently observed in *Papilio* (*29*). To test for the genetic basis of the two rules in this system, we carry out QTL studies of body size (pupal weight) and female reproduction (ovary dysgenesis) in backcross hybrids. We then compare *Papilio* with *Heliconius* (*26*) to test whether these genera share a similar genomic basis for the two rules.

**Figure 1.**
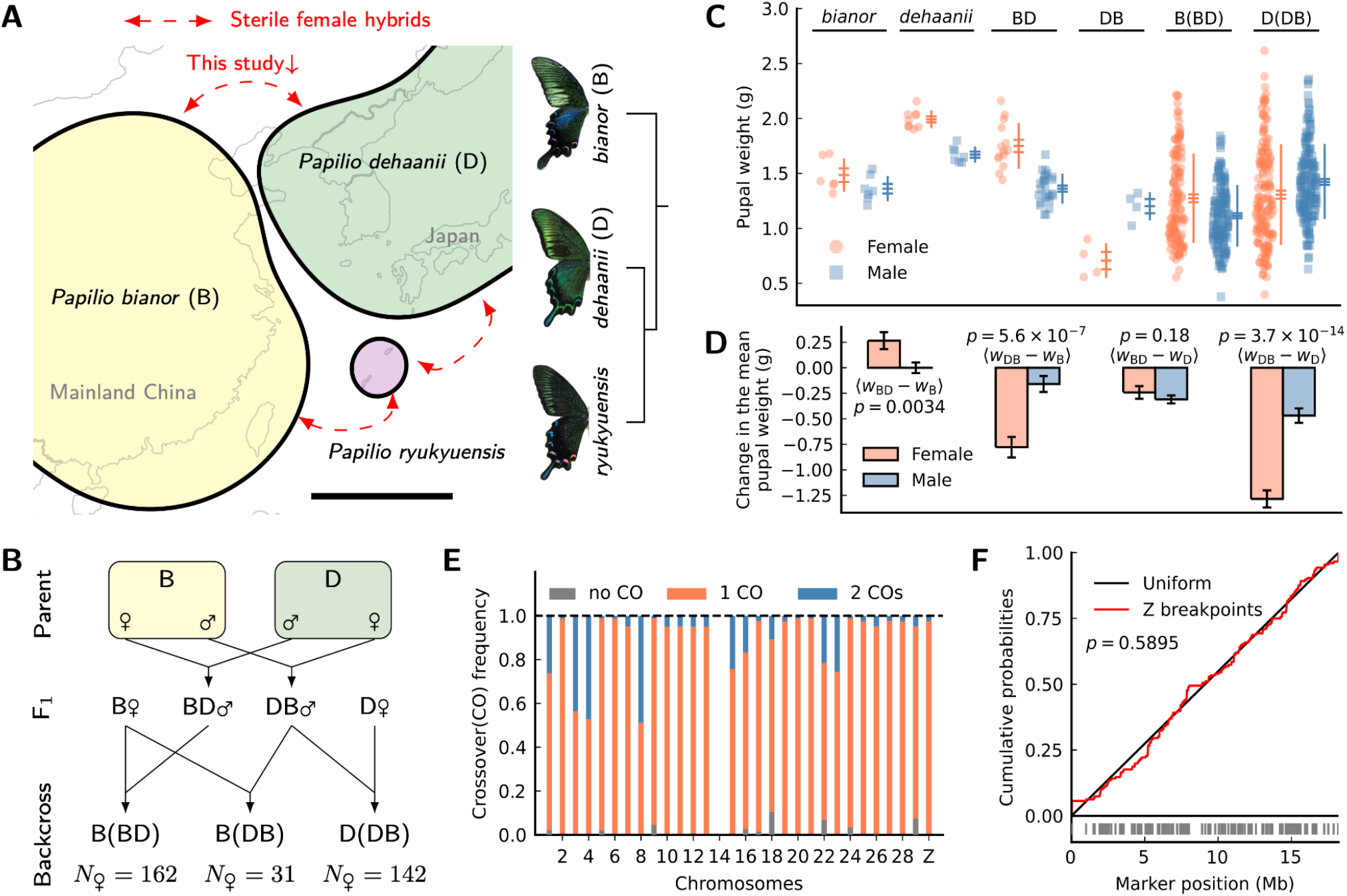
The study design, variation in pupal weight, and patterns of meiotic crossover in males. **(A)** Geographic distribution and species relationships in the *Papilio bianor* complex. Scale bar=1000km. **(B)** Crossing design. **(C)** Pupal weight variation among parents, F_1_s, and backcrosses. Horizontal bars represent the mean +/-standard error. Vertical lines represent the mean +/-standard deviation. **(D)** Changes in mean pupal weight between F_1_s and parents in each sex. Error bars are standard errors of the mean changes. The significance of male vs female differences is shown as p-values of Z-tests. **(E)** The crossover frequency in F_1_ males per chromosome pair per meiosis. Chr14 is excluded due to assembly problems. **(F)** Recombination breakpoints on the Z chromosome are uniformly distributed (p-value is from a Kolmogorov-Smirnov test). Vertical bars are inferred breakpoints.

### Haldane’s Rule between *P. bianor* and *P. dehaanii* involves asymmetrically inherited elements

To investigate hybrid abnormality, we performed reciprocal F_1_ crosses and backcrosses between the two species (Fig. 1B). We follow the order (female × male) in notation. For instance, “B(BD)” is equivalent to “*bianor* ♀ × (*bianor* ♀ × *dehaanii* ♂) ♂”, where “B” and “D” stand for *bianor* and *dehaanii*, respectively.

Pupal weight (*W*) is treated as a proxy for adult body size. F_1_ females with a *dehaanii* mother (“DB”) are significantly smaller than females of either parental species, but in the reciprocal cross with a *bianor* mother (“BD”) they span the range of parental females (Fig. 1C). For F_1_ males, deviation in pupal weight from pure males is less extreme than that of F_1_ females (Fig. 1D). We interpret female-biased abnormal size as a defect conforming to Haldane’s Rule.

Focusing on hybrid female sterility, we dissected ovaries across the pedigree and determined major ovary phenotypes (Fig. 2). To our surprise, while F_1_ females with a *bianor* mother (“BD”) have almost empty ovaries (Fig. 2C, 2J), F_1_ females in the reciprocal cross (“DB”) develop and lay superficially normal eggs (Fig. 2B, 2I). However, eggs laid by mated females never hatch. We score ovary phenotype rather than female fertility *per se*: ovaries with regularly spaced and spherical follicles are subsequently all classified as “Normal”. Since deformed ovaries lead to sterility and can be readily scored, this is a sufficient and efficient approach to detect sterility factors.

**Figure 2.**
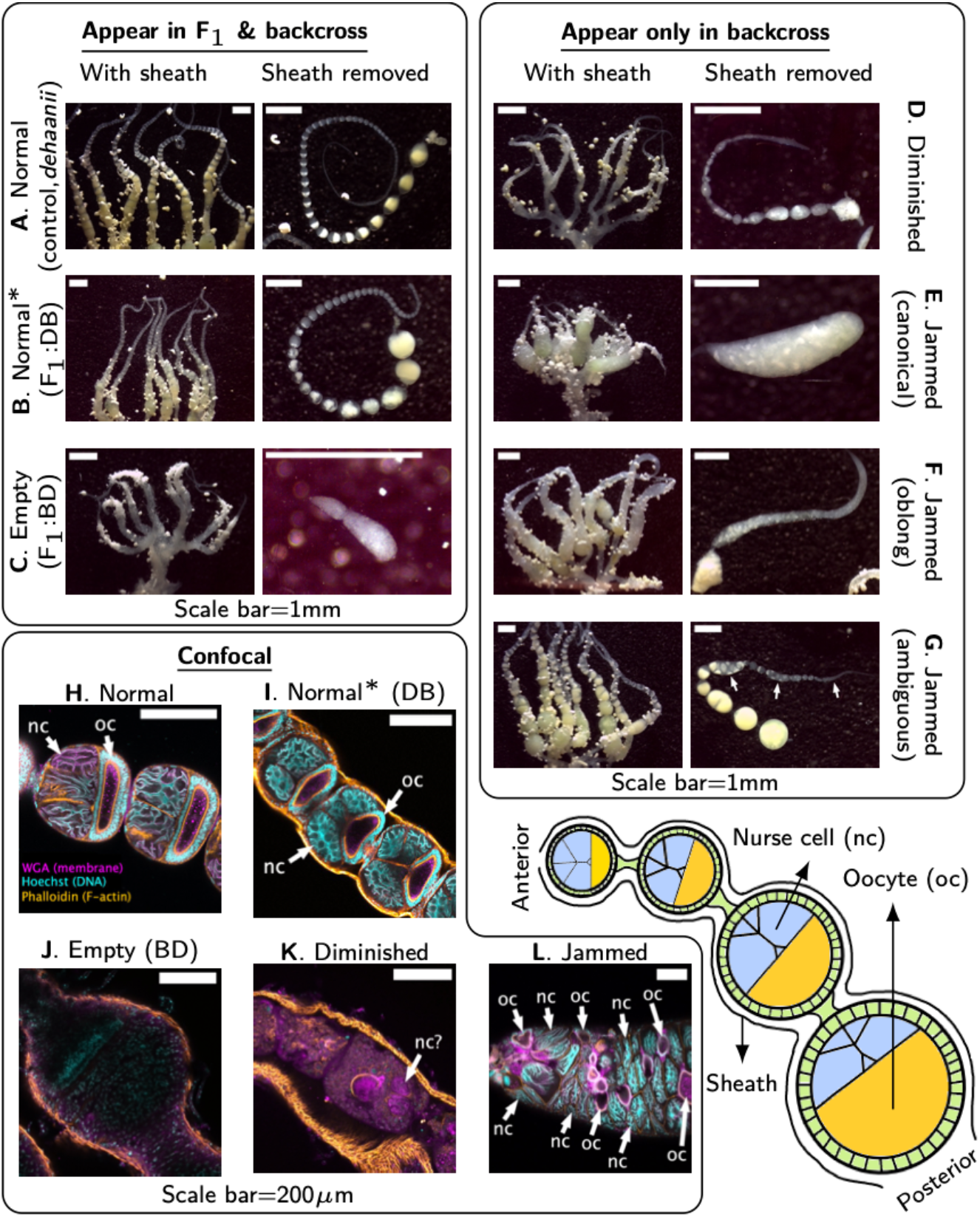
Ovary morphology supports a categorical classification of phenotypic defects in *Papilio.* Monochrome confocal images for each channel are in Fig. S1 and S2 (WGA stains membranes in magenta; Hoechst stains DNA in cyan; Phalloidin stains F-actin in orange). A schematic diagram of butterfly ovarioles is shown at the bottom right. Scale bars for stereoscope images are approximate. **(A, H)** Phenotype Normal in a pure individual. The confocal image shows follicles with sheath removed. **(B, I)** Phenotype Normal in F_1_ females of the cross DB. Sheath is retained in the confocal image. **(C, J)** Phenotype Empty in F_1_ females of the cross BD. Sheath is retained in the confocal image with little tissue inside. **(D, K)** Phenotype Diminished. Sheath is retained in the confocal image with a substantial amount of interior tissue without discernible follicle structures. **(E-G, L)** Phenotype Jammed. Sheath is removed in the confocal image, and follicle cells fuse into tubes with many nurse cells and oocytes. Subtype canonical: each ovariole collapses into a single tube of oocytes and nurse cells. Subtype oblong: tubes are elongated. Subtype ambiguous: tubes and isolated follicles coexist.

Ovary and female size phenotypes differ significantly between reciprocal crosses and are therefore examples of “Darwin’s corollary” of hybrid incompatibility (*30*). This phenomenon is likely due to asymmetrically inherited genetic elements. For this reason, we separate backcross types according to maternal origin in QTL analysis.

### Patterns of meiotic crossover on the Z chromosome in F_1_males

Understanding how recombination determines chromosomal ancestry is crucial to our subsequent analysis. To infer haplotypes and crossover patterns, we carried out whole-genome low-coverage (∼1x) sequencing in backcrosses, while F_1_s and parents were sequenced to higher depths (>5x and >30x, respectively). Prior to crossover analysis, we used linkage information from all families to correct assembly errors in the reference genome of *P. bianor* (*31*), except for chromosome 14, where errors remained unresolved (Fig. S4). As assembly errors can affect inference of recombination breakpoints, we do not report crossover patterns on chromosome 14.

We inferred the crossover pattern in F_1_ males by counting the estimated recombination breakpoints across all backcross offspring (female meiosis in Lepidoptera lacks crossovers). Most F_1_ males had at least one crossover per chromosome pair per meiosis, but the degree of crossover interference varied among chromosomes (Fig. 1E). Double crossovers were frequent on some chromosomes (>40% on chromosome 8) but very rare on most. Importantly, the Z chromosome in the *Papilio* crosses had almost no double crossovers, and its recombination breakpoints were approximately uniformly distributed along the chromosomal axis (Fig. 1F). Recombination on the Z chromosome can thus be approximated by a model where the first crossover occurs uniformly and randomly across the chromosome, while the second crossover is completely suppressed. We apply this model to the study of incompatibilities.

### Abnormal pupal weight is determined by multiple additive factors scattered across the Z chromosome

Pupal weight in offspring of B(BD) and D(DB) backcrosses has a broad distribution (Fig.1C). Such a distribution could be associated with multiple chromosomes or extreme developmental stochasticity. Yet single-marker QTL scans suggest that the Z chromosome alone controls pupal weight variation in females (Fig. 3A). These scans reveal a major QTL near the center of the Z chromosome in both cross types, explaining over 50% of the phenotypic variance (Fig. S8). Nonetheless, we reason below that this major QTL is likely a statistical artifact caused by multiple factors scattered across the Z chromosome.

**Figure 3.**
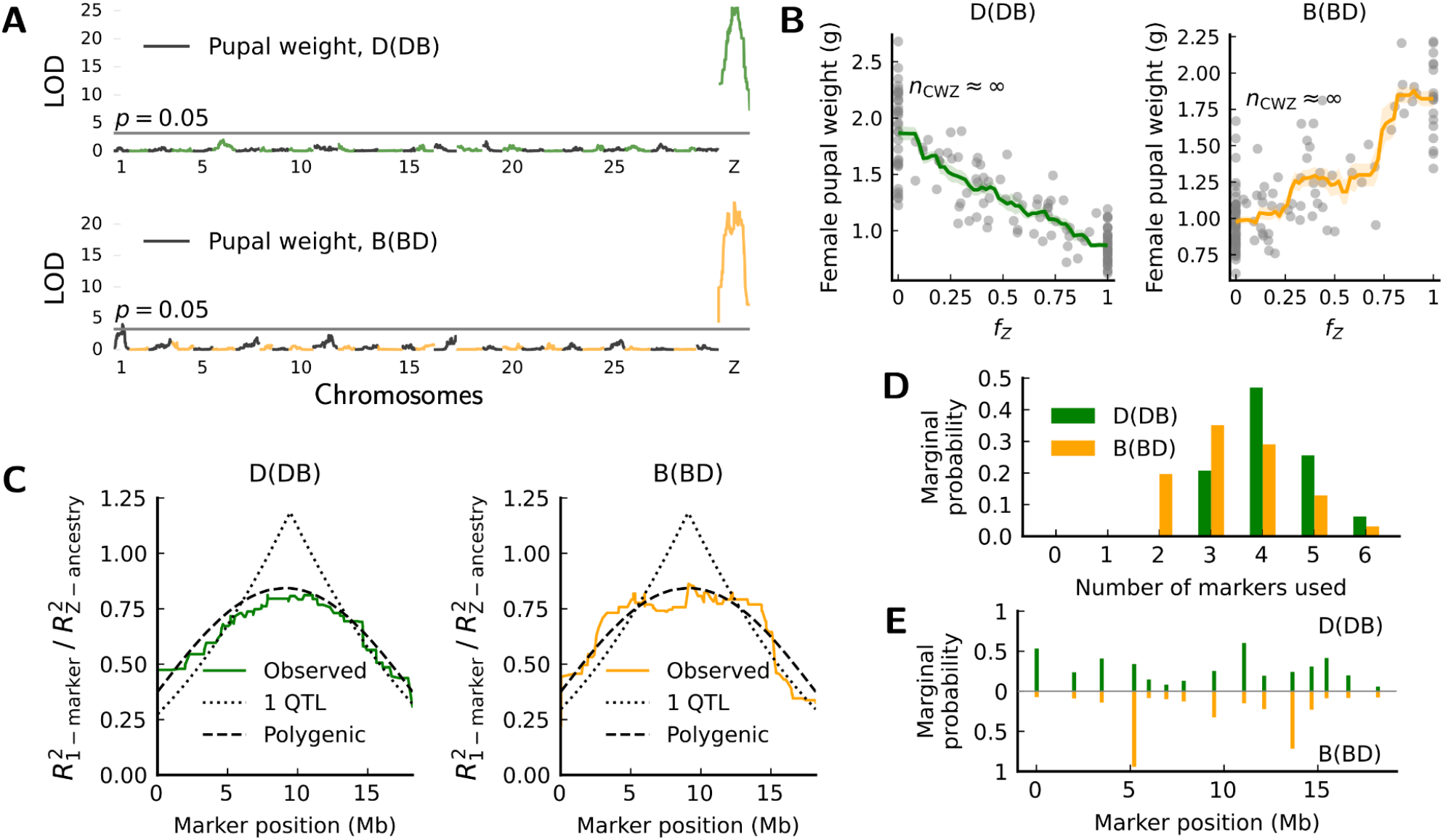
Pupal weight variation in female hybrids maps to many additive Z-linked effects. **(A)** One-dimensional QTL scan shows that variation in pupal weight is mainly explained by the Z chromosome. LOD scores peak at the Z chromosome center. Alternating colors represent different chromosomes. **(B)** Ancestry fraction on the Z chromosome (*f*_*Z*_) is highly informative of pupal weight. A crude estimator of the effective number of genes (*n*_CWZ_) (*32*) using segregating phenotypic variance approaches infinity. **(C)** Genotype-phenotype regression does not support any 1-QTL architecture for pupal weight. The distribution of regression power is and *R*^*2*^_1-marker_/*R*^*2*^ _Z-ancestry_ (the ratio between *R*^*2*^ using a single marker versus a “polygenic” model of Z chromosome ancestry). *R*^*2*^_1-QTL_ and *R*^*2*^ _polygenic_ are the fit between observed and theoretical curves. The polygenic model provides a good fit for the observed curves. **(D, E)** Bayesian QTL model selection (*33*) based on 15 sparsely spaced markers favors the simultaneous inclusion of multiple markers (≥3) in predicting pupal weight. Markers more favored by models have higher marginal probabilities of being selected. Marginal probabilities equal the partial sum of the posterior probabilities of QTL models.

The following evidence supports Z-linked polygenic architecture. First, pupal weight depends almost continuously on the fraction of introgressed ancestry on the Z chromosome (Fig. 3B; from here on the Z-chromosome ancestry fraction is denoted as *f*_*Z*_, and the autosomal ancestry fraction as *f*_*A*_). Second, genetic variance of pupal weight is much smaller than that expected for a single locus of major effect, so that a crude estimate of the number of loci (*n*_CWZ_) (*32*) using segregating genetic variance is effectively infinite (Table S1). Thus, multiple additive factors may be involved. Third, we compare two extreme genetic architectures: a single-QTL model versus a polygenic additive model in which weight varies linearly with *f*_*Z*_. Given the polygenic model, the strength of the genotype-phenotype association (*R*^*2*^) will always be larger for models using *f*_*Z*_ (i.e., *R*^*2*^_1-marker_/*R*^*2*^_Z-ancestry_<1 for all markers). Conversely, given the single-QTL model, markers tightly linked to the QTL will surpass *f*_*Z*_ in association strength (i.e., *R*^*2*^_1-marker_/*R*^*2*^_Z-ancestry_>1 for some markers). Using the approximate crossover model, we derive *R*^*2*^_1-marker_/*R*^*2*^_Z-ancestry_ on the Z chromosome analytically under each architecture (Supplementary Text–2.2), and the observed patterns closely resemble polygenic predictions (Fig. 3C). Fourth, Bayesian QTL model selection (*33*) also favors multiple additive markers in predicting pupal weight (Fig. 3D).

Posterior probabilities for markers being selected are more evenly distributed in D(DB) females (Fig. 3E, top), congruent with a near-linear relationship between pupal weight and *f*_*Z*_ (Fig. 3B, left). For B(BD) females, this relationship is less smooth (Fig. 3B, right), and a two-QTL model is a slightly better fit to the observed *R*^*2*^_1-marker_/*R*^*2*^_Z-ancestry_ curve (Fig. S9). Nonetheless, the two apparent QTLs flank the chromosome center symmetrically with similar effects, and this is indistinguishable from a polygenic model with abrupt weight increase at two *f*_*Z*_ thresholds (*34*). Finally, the polygenic model offers a simple explanation for the single QTL at the Z chromosome center: after a single crossover, central markers are individually more informative of *f*_*Z*_ than markers near chromosome ends (Supplementary Text: Theorem 2). Central markers thus provide richer phenotypic information when phenotype varies almost linearly with *f*_*Z*_, generating a statistical peak in association strength. In summary, while the exact number of Z-linked factors remains unknown, our evidence is highly consistent with a multigenic architecture, rendering the Z-chromosome ancestry fraction informative in predicting phenotype.

### Polygenic basis of ovary dysgenesis on the Z chromosome

Ovary dysgenesis leads to hybrid female sterility. The only previous attempt to map hybrid sterility factors genome-wide in Lepidoptera found a pair of epistatic QTLs near each end of the Z chromosome in the backcross from *Heliconius pardalinus sergestus* to *H. p. butleri* (Fig. 4A, left) (*26*). Additionally, that study suggested a weak single-locus QTL near the Z chromosome center (Fig. 4B, top). In *Papilio* D(DB) females, two-marker scans also identified a pair of epistatic QTLs on the Z chromosome for phenotype “Normal” (Fig. 4A, right), although we found no single-locus QTL on the Z chromosome (Fig. 4B, bottom).

**Figure 4.**
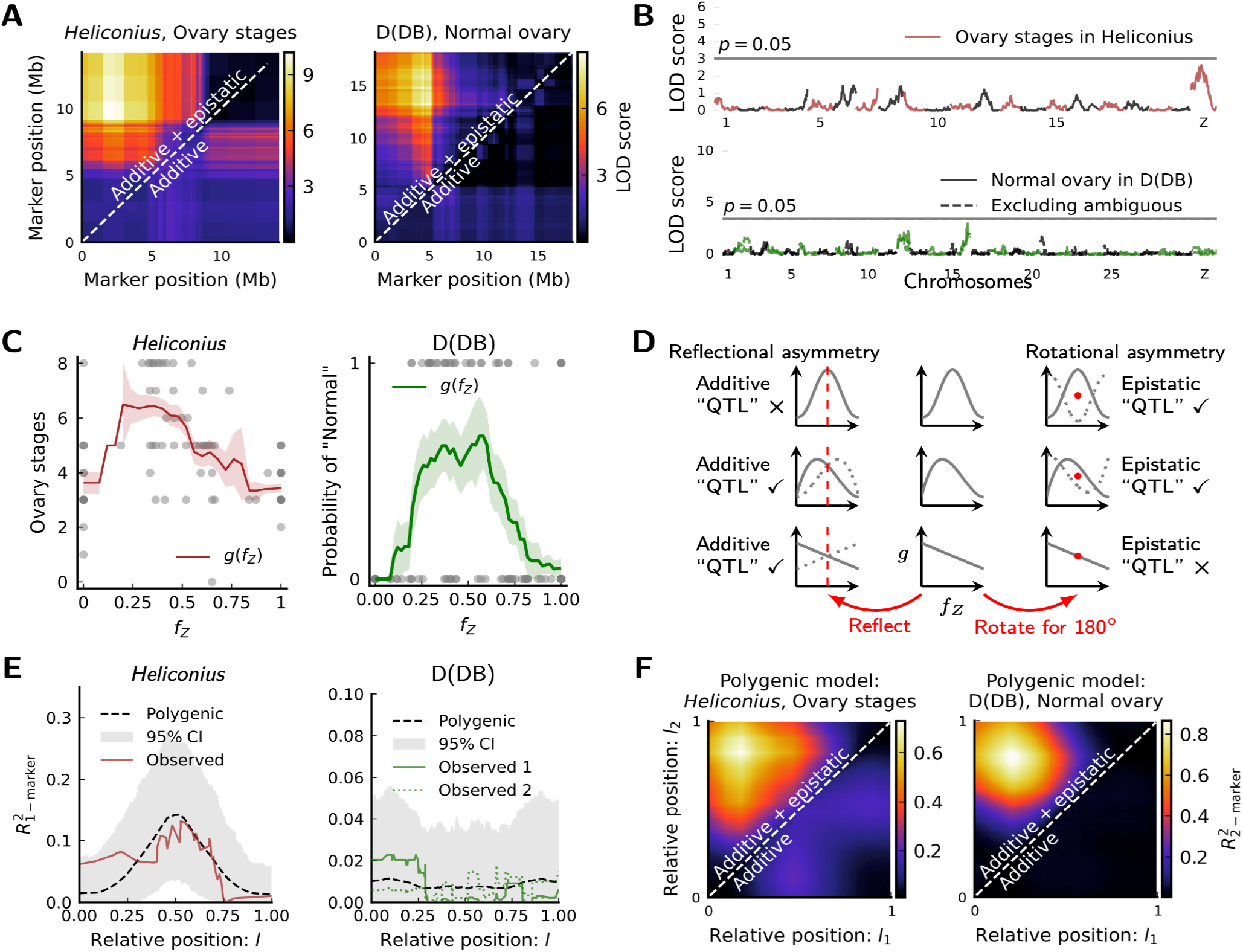
The polygenic basis of ovary dysgenesis in *Heliconius* and D(DB) females. **(A)** Two-marker scans identify a pair of epistatic QTL located near both ends of the Z chromosome. **(B)** One-marker scans identify a very weak additive QTL on the Z chromosome in *Heliconius* (previously reported to be significant by a different scoring method). No significant additive QTL on the Z chromosome in D(DB) females. **(C)** Ancestry fraction on the Z chromosome (*f*_*Z*_) is informative of ovary phenotype. The moving average of phenotype scores (window size=0.2) is used as the polygenic model *g*(*f*_*Z*_). Shaded areas are means +/-standard errors. Dots in D(DB) represent one round of phenotype assignment due to ambiguous individuals. **(D)** If a trait is fully polygenic on the Z chromosome, QTL analysis via marker-phenotype regression only reveals the asymmetry in *g*(*f*_*Z*_). The presence of additive “QTL” is equivalent to a reflectional asymmetry in *g*, while epistatic “QTL” is equivalent to a rotational asymmetry in *g*. These conclusions are valid for the inferred Z-chromosome crossover model. **(E)** Polygenic models recover observed results of one-marker scans. Dashed curves and shaded areas represent the means and 95% confidence intervals of *R*^2^_1-marker_, across 1,000 independent draws of individuals matching real sample sizes (*N*_*Heliconius*_=86, *N*_D(DB)_=142). For D(DB), observed 1 classifies ambiguous individuals as “Normal”, while observed 2 classifies them as the opposite. **(F)** Polygenic models recover observed epistasis in two-marker scans. Heatmaps show the means of *R*^2^ _2-marker_ across 100 independent draws of individuals matching real sample sizes.

Backcross females in *Heliconius* and D(DB) females in *Papilio* develop more normal ovaries when the Z chromosome is recombined, regardless of the precise chromosomal location of introgressed ancestry (Fig. 4C, S11). Since the Z-chromosome ancestry fraction carries so much phenotypic information, we argue that the architecture may again be polygenic on the Z chromosome for both cases, and that the mapped epistatic QTLs on the Z chromosome are likely artifacts. We extend the polygenic model developed previously for pupal weight to ovary dysgenesis. We hypothesize that the expected ovary phenotype is in general a continuous function (*g*) of the Z-chromosome ancestry fraction (*f*_*Z*_). In practice, *g* is the moving average of phenotypic scores with respect to *f*_*Z*_ (Fig. 4C). Again, if phenotypes depend only on ancestry fraction, a significant QTL does not necessarily imply a major effect of the identified locus in development. Rather, these QTLs arise indirectly from the asymmetry in *g* (Fig. 4D). With the inferred crossover model, we prove for backcrosses that:

i. In one-marker scans, additive QTLs are caused by the reflectional asymmetry in *g*(*f*_*Z*_) with respect to its center (Fig. 4D left. See Supplementary Text: Theorems 6-8 and Fig. S12 for details)
ii. In two-marker scans, epistatic QTL pairs are caused by the rotational asymmetry in *g*(*f*_*Z*_) with respect to its center (Fig. 4D right. See Supplementary Text: Theorems 9, 10 and Fig. S14 for details)

For both cases of ovary dysgenesis, *g* has a unimodal form (Fig. 4C). This form is strong in rotational asymmetry but weak in reflectional asymmetry (e.g., the first two rows in Fig. 4D). Consequently, it predicts strongly epistatic QTLs but no or only weak additive QTL, as observed (Fig. 4E, 4F). In contrast, the *g* for pupal weight is largely linear, which is only reflectionally asymmetric (e.g., the last row in Fig. 4D). This shape predicts a major additive QTL at the chromosome center (Fig. 3C) but no epistatic QTLs (Fig. S10). Thus, not only does the Z-chromosome ancestry fraction carry phenotypic information, but it also predicts the strength of marker-phenotype association across the Z chromosome, and can explain the presence of both additive and epistatic apparent QTLs. This reasoning therefore supports a polygenic basis of the large-Z effect in both systems.

### Modulation of incompatibilities by autosomal and maternal backgrounds

Autosomal and maternal genetic backgrounds are also important in modulating hybrid defects. For pupal weight, if Z-linked additive effects were independent of genetic background, F_1_ females between a *dehaanii* mother and a *bianor* father should be of similar size as the paternal species. Instead, they are much smaller than either parental female (Fig. 1C), implying epistasis between the *bianor* Z chromosome and the *dehaanii* background. Likewise, pupal weight in B(BD) females is much smaller than both parental species when *bianor*’s Z chromosome contains no introgression (Fig. 3B). This stable decrease in pupal weight can only be explained by the ∼25% autosomal introgression from *dehaanii*, implying again that such Z-linked effects could arise via interactions with autosomes, albeit in the opposite maternal background. Similarly, ovaries are defective in *Heliconius* as well as *Papilio* D(DB) backcrosses even when the Z chromosome comes entirely from the maternal species (Fig. 4C), consistent with a significant role of autosomal introgression in defect development. Defects induced via autosome-only introgression develop rather stably (Fig. 3B, S11C, S13A), indicating that multiple autosomal factors linked loosely are necessary to prevent phenotypic segregation. These results suggest that autosomal components of incompatibility are also somewhat polygenic, and may depend on the autosomal ancestry fraction (*56*).

### Exceptions to polygenic architecture

The polygenic architecture we have found, however, does not apply to all incompatibilities in *Papilio*. In females with a *bianor* mother, introgression of a small region (∼1Mb) on the Z chromosome from *dehaanii* is sufficient to induce the defective ovary phenotype “Empty” (Fig. S15A, Table S2). This phenotype obliterates nearly all follicle tissues in ovarioles (Fig. 2C, 2J), effectively blocking all other ovary defects. In this maternal background, a small region on chromosome 8 also modulates the development of the “Normal” ovary phenotype (Fig. S15B, Table S2). These results suggest that ovary dysgenesis in *Papilio* from this direction of cross is predominantly affected by narrow genomic regions of large effect, in contrast to the polygenic architecture in the opposite maternal background.

## Discussion

### Genetic logic underlying two rules of speciation

Dominance theory requires incompatibility factors to be recessive on the sex chromosome so that the homogametic sex is sheltered by dominance (*7, 14*). This theory is applicable to Lepidoptera but faces obstacles here. First, polygenicity implies that a large number of Z-linked incompatibility factors should be simultaneously recessive, which is untested in Lepidoptera. Second, intermediate levels of introgression on the Z chromosome rescue ovary defects in backcross females (Fig. 4C). This is unexpected under dominance theory because intermediate levels of introgression would still expose recessive Z-linked factors in females. Faster-Z theory has little support either: the *dN/dS* ratio is similar between the Z chromosome and autosomes in *Heliconius* (*26*), and sequence divergence in *Papilio* is even greater on autosomes (Fig. S16).

Polygenic incompatibilities are, however, congruent with mechanisms based on asymmetric inheritance (*30*). Recall that *f*_*Z*_ and *f*_*A*_ are the introgressed ancestry fractions on the Z chromosome and on all autosomes, respectively. The key insights are:

i. *f*_*Z*_ is more skewed from *f*_*A*_ in F_1_ females than in F_1_ males;
ii. *f*_*Z*_ is much more variable than *f*_*A*_ in backcrosses;
iii. *f*_*A*_ is reduced in backcrosses compared to that in F_1_ hybrids.

Since intermediate *f*_*Z*_ produces more normal phenotypes in backcrosses, a balancing process appears to exist between ancestry on autosomes and on the Z chromosome: phenotypes degrade when *f*_*Z*_ and *f*_*A*_ deviate from optimal balance. This process is ancestry-based and requires a multilocus genetic basis.

Additionally, i) implies that F_1_ females are more extremely unbalanced, generating Haldane’s Rule; ii) implies that variation of imbalance is largely attributable to the Z chromosome, generating a large-Z effect; and iii) implies that the optimal Z-chromosome ancestry fraction might differ between F_1_ and backcrosses.

### Interpreting QTL results of highly polygenic traits

We find that QTL analysis of highly polygenic traits is shaped strongly by the statistical structure of ancestry tracts. Spurious major effect loci may appear in association mapping using crosses with long ancestry tracts even with large sample sizes. Here, “ghost” QTLs are likely to result from the cumulative influence of all polygenes linked to focal markers, and a peak in association strength does not result from a major single locus effect in development. This problem has been well recognized in genetic association theories despite a lack of universal solutions (*34*–*37*). Still, empirical analyses rarely consider this complexity because common software assumes only one or two QTLs per chromosome (*38, 39*). The remedy we adopt here is to model explicitly the generating process of ancestry tracts (e.g., a crossover model), and to integrate information across all markers in picking the best-fitting architecture. Polygenic traits are common in humans (*40*), and between-species traits such as hybrid incompatibilities can be more complex due to greater genomic divergence. A stronger test for polygenicity needs to incorporate many more configurations of ancestry blocks than available in our backcrosses.

### Polygenic incompatibility and global epistasis

Hybrid incompatibility is usually perceived as negative epistasis between species-specific mutations (*41, 42*). While our phenomenological explanation invokes epistasis, the interaction may well be between ancestry fractions in different genomic regions. This is similar to global epistasis, where the phenotypic effect of a genetic change is somewhat independent of specific loci underlying the change (*43*). Global epistasis can emerge as a transformation of additive components. In our case, global epistasis is manifested as *g* (transformation) on ancestry fraction (additive components). Global epistasis has been found previously in polygenic incompatibility. For instance, hybrid male sterility in *Drosophila* is sensitive not to precise genomic regions but to the total introgressed ancestry fraction (*44*– *46*). The molecular nature of polygenicity is unresolved. In our case, it is tempting to consider epigenetic mechanisms between autosomes and the Z chromosome. For instance, genetic variance of pupal weight in backcross males is much smaller than in females (Table S3). This is consistent, for instance, with dosage compensation in Lepidoptera in which both Z chromosomes in males are partially suppressed (*47*), which will dampen the effects of introgressed alleles. When we embarked on these studies of hybrid incompatibility in Lepidoptera, we hoped to locate key genes causing defects. To our surprise, it seems clear instead that sterility and other incompatibilities are often polygenic, and that this polygenicity can provide simple explanations for some hitherto mysterious rules of hybrid incompatibility.

## Acknowledgments

We thank Naomi Pierce and Adam Cotton for providing background information on the study system; Janet Sherwood, Shui Xu, Yuchen Zheng, Jinbo Hu, and Anastasios Kougionis for their assistance and knowledge in breeding/sourcing host plants and butterflies; Cassandra Extavour for discussing oogenesis; John Wakeley, Robin Hopkins, Liang Qiao, Sarah Dendy, Nathaniel Edelman, Shuzhe Guan, Fernando Seixas, and Yuttapong Thawornwattana for their intellectual support during the project.

## Funding

The Quantitative Biology Initiatives at Harvard University (TX); The National Science Foundation-Simons Center for Mathematical and Statistical Analysis of Biology at Harvard University #1764269 (TX); The Department of Organismic and Evolutionary at Harvard University (TX, ST, JM); Sigma Xi Grant in Aid of Research (TX); Harvard University GSAS Student Council Summer Research Grant (TX)

## Author contributions

Conceptualization: TX, JM; Methodology: TX, JM; Software: TX, NR; Formal analysis: TX; Investigation: TX, ST; Resources: XL, MY; Data curation: TX; Visualization: TX, ST; Funding acquisition: TX, JM; Project administration: TX; Supervision: JM; Writing – original draft: TX, ST, JM; Writing – review & editing: TX, ST, NR, XL, MY, JM

## Competing interests

The authors declare no competing interests

## Data and materials availability

Raw reads are released in the NCBI Sequence Read Archive (BioProject: PRJNA892033). Source data and code for main and supplementary figures will be deposited in Dryad and will be released at the time of publication. Source code is also available from: https://github.com/tzxiong/2022_Papilio_HybridIncompatibilityMapping

## Notes

https://github.com/tzxiong/2022_Papilio_HybridIncompatibilityMapping

### Competing Interest Statement

The authors have declared no competing interest.

### Summary of Updates

Major revision to the QTL results. Incorporated more analysis of Heliconius. Derived mathematical theorems for the existence of "ghost" QTLs. Clarifies why the proposed mechanisms is distinct from existing theories.

