## Supplementary Materials for "Polygenic Effects on the Z Chromosome Underlie Hybrid Incompatibility in *Papilio* and *Heliconius* Butterflies"

### 1. Materials and Methods

#### 1.1 Breeding

Lineages of *P. dehaanii* were purchased directly from a butterfly farm in Qingdao (Shandong Province, China), exclusively sourced from a small local population. Lineages of *P. bianor* were collected in the field from Ningbo (Zhejiang Province, China) for breeding in 2020, 2021. A few individuals were also collected in the field from Kunming (Yunnan Province, China) for breeding in 2019. All crosses were done by hand-pairing. Eggs were collected by putting females in small cages with host plants under fluorescent light. The following host plants were used throughout the project: *Tetradium daniellii*, *Zanthoxylum bungeanum*, *Z. ailanthoides*, *Z. beecheyanum*, *Z. simulans*, *Choisya ternata*, *Phellodendron amurense*. Larvae and pupae were kept in greenhouse conditions (approximately 20°C∼35°C), with a combination of natural and greenhouse lights to maintain at least 10 hours of illumination per day. Adults for dissection were immediately put into a 5°C room after eclosion to reduce activity. Otherwise, they were fed with sugar water once a day, and females were subsequently kept in the dark, while males were in an illuminated growth chamber to facilitate hand-pairing.

#### 1.2 Phenotyping ovaries

Ovaries were dissected from females within five days of eclosion in 1× PBS solution. Ovariole sheath was manually removed, and most images were taken using the internal camera of a Leica EZ4 HD stereo microscope (pictures of a few specimens were taken by a cellphone through the eyepiece of a Zeiss Stemi 2000 stereo microscope). Due to a limitation of the stereoscope, many stereoscope images were not scaled exactly at the time of image acquisition. The approximate scales of these images were determined by comparing magnification levels against images with pre-determined scales. This level of inaccuracy does not affect phenotypic scores because only qualitative differences (i.e., ovariole shapes and the presence/absence of certain structures) were used to classify phenotypes. Since ovary phenotypes are categorical, we established all major categories by defining the most obvious and the most frequent phenotypes across all dissected ovaries. These categories were confirmed later by confocal imaging that they have significant qualitative differences (see Fig. 2 in the main text). Phenotype Jammed is variable in terms of the fraction and the position of Jammed follicles. We lumped variable forms of Jammed into a single category in QTL analyses to reduce human bias in separating different kinds of Jammed.

A small number of ovaries have ambiguous phenotypes for one of the following reasons: 1) Different ovarioles develop different phenotypes; 2) Some part of the ovary is lost in dissection; 3) Extremely rare phenotypes resembling none of the existing categories. These ambiguous individuals, mostly from D(DB) females, were assigned multiple categories. When such uncertainty affects analyses, two methods were used: 1) For simultaneous analysis of more than two ovary phenotypes: randomly select a phenotype from previously assigned categories on ambiguous individuals, perform analyses, and repeat the same procedure many times; 2) For QTL mapping in software r/qtl or r/qtl2 with binary categorical traits: map QTLs with ambiguous individuals scored as 0 (and 1) in the first (and the second) attempt.

#### 1.3 Staining and confocal imaging of ovaries

Dissected ovaries were fixed in 4% Paraformaldehyde solution in 1× PBS for 20 minutes at room temperature. The ovaries were washed for 15 minutes each in 0.1% PBTx ( 1× PBS, 0.1% Triton-X 100), 1% PBTx, 2% PBTx, 0.01% Saponin (Sigma Aldrich 47036) in 1× PBS and Blocking solution (1X PBS, 0.3% Triton-X 100, 0.5% Normal Goat Serum). The ovaries were then stained for 12 hours using the following reagents at 1:500 dilution in blocking solution: Hoechst 33342 (10 mg/ml, Thermo Fisher H3570), Wheat Germ Agglutinin-647 (WGA, Thermo Fisher W32466) and Rhodamine Phalloidin (Thermo Fisher R415). Stained ovaries were washed four times for 15 minutes each in 0.1% PBTx followed by a final 1× PBS wash. After the washes, ovaries were mounted in equal volumes of 1× PBS and Vectashield mountant (Vector labs H1900) on a slide.

The ovary samples were imaged by acquiring Z-section images on a Zeiss LSM 880 laser scanning confocal microscope at Harvard Center for Biological Imaging. The microscope was equipped with an Argon laser and a He/Ne 633 nm laser. Zeiss Plan-Apochromat 10×/0.45 M27 or 20×/0.8 M27 objective lenses were used for imaging. All images were 1024 × 1024 pixels in size and were acquired using PMT detectors. Images were acquired at excitation/emission wavelengths of 405/450 nm for Hoechst 33342, 561/610 nm for Rhodamine Phalloidin, and 633/696 nm for WGA.

#### 1.4 DNA extraction and sequencing

Samples from the cross were preserved in either pure ethanol or RNAlater at -20°C prior to DNA extraction. For extraction, we used E.Z.N.A Tissue DNA kits (Omega Bio-tek, Inc.). Whole-genome library preparation was performed using Illumina DNA 1/4 reactions kits at Harvard University Bauer Core, barcoded, and subsequently sequenced altogether on a single lane of Illumina NovaSeq S4. Autosomal coverage varies among individuals: backcrosses-1×, $F_1$s-5×, parents-30× to 60×. Raw reads were trimmed with Cutadapt-3.4 (*48*) to remove adapters (CTGTCTCTTATACACATCT), and subsequently mapped to the reference genome of *P. bianor* using the BWA-0.7.17 MEM algorithm. Duplicate reads were marked using Picard-2.25.7 (*49*). We used BCFtools-1.9 (*50*) to pile up reads with very light quality filtering, and called variants with associated genotype likelihoods. VCF files produced by the variants caller were used for linkage analysis.

#### 1.5 Linkage analysis

For quality control, we first calculated kinship coefficients among individuals using NgsRelate-2 (*51*) and corrected the pedigree position of a few individuals (Fig. S3). Lep-MAP-3 was used for all subsequent linkage analysis (*52*). First, VCF files and the pedigree were combined in module ParentCall2, and we imputed haplotype structure along the reference genome with module OrderMarkers2. We also generated de novo marker orders using the same module. Comparing de novo marker orders against the order in the original reference genome revealed some intra-chromosome assembly problems. We corrected these problems below.

#### 1.6 Linkage-based reference genome correction

For a pre-correction reference genome (31), we plotted the de novo marker order against the genomic order. We discovered some reference assembly errors that affect the order and orientation among PacBio scaffolds on chromosomes. Using inferred haplotypes in the "grandparental" phase (i.e., the parents from the cross. This terminology "grandparental phase" is used in Lep-MAP-3), we calculated the correlation of ancestry between each pair of markers, which should be a decreasing function of marker distance on each chromosome due to recombination. This information enabled us to correct large-scale errors and an intermediate reference genome was generated. To correct smaller errors, we inferred de novo marker order by Lep-MAP-3 on the intermediate genome and compared it against the intermediate genome. This extra step corrected the position/orientation of several smaller scaffolds. We were able to correct all errors discovered in this way except for those on chromosome 14, where an apparent orientation problem appears to occur *within* a PacBio scaffold and we were unable to determine its breakpoint. (This may be due to a different population used here compared to the reference population). The de novo marker order inferred from this final genome is mostly collinear to the genomic order (Fig. S4), confirming that most visible errors in concatenating scaffolds have been eliminated. Finally, haplotypes in the grandparent phase were re-inferred using the corrected reference genome for all subsequent analyses.

#### 1.7 Inferring crossover frequency

To infer crossover frequency, we counted the number of recombination breakpoints in each paternal haplotype among all backcross individuals (For all paternal haplotypes, see Fig. S5, S6, S7). No chromosome has more than two breakpoints except for chromosome 14 (likely due to the aforementioned reference error). We excluded chromosome 14 from this crossover analysis. Let $n_0$, $n_1$, and $n_2$ be the number of haplotypes having 0, 1, or 2 recombination breakpoints for a given chromosome. The maximum likelihood estimate of crossover frequency is as follows (see "Supplementary Text-2.1" for derivation). First, calculate:

$$
\begin{aligned}
c_0 &= (n_0 - n_1 + n_2)/(n_0 + n_1 + n_2) \\
c_1 &= (2n_1 - 4n_2)/(n_0 + n_1 + n_2) \\
c_2 &= 4n_2/(n_0 + n_1 + n_2)
\end{aligned}
\tag{S1}
$$

If $c_0, c_1, c_2$ are all nonnegative, they are the inferred frequencies of having 0, 1, or 2 crossovers. If $c_0 < 0$, the adjusted estimate is

$$
\begin{aligned}
c_0^* &= 0 \\
c_1^* &= (n_0 - n_2)/(n_0 + n_2) \\
c_2^* &= 2n_2/(n_0 + n_2)
\end{aligned}
\tag{S2}
$$

#### 1.8 QTL scans by marker regression in r/qtl and r/qtl2

One-dimensional QTL scans were performed with R-package qtl2 (*39*), and two-dimensional scans were performed with R-package qtl (*38*). To calculate LOD scores, ovary phenotypes were always mapped one at a time using a binary trait logistic mapper (i.e., phenotype of interest = 1, other phenotypes = 0). This approach is suitable for unordered categorical traits such as ovary morphology[1]. For pupal weight, we introduced brood as a covariate to control for seasonal variation in pupal weight due to diet and environmental factors. LOD score thresholds were always estimated on 1,000 random permutations of phenotypes.

#### 1.9 The number of loci underlying a quantitative trait using segregating genetic variance

A crude way to test if multiple loci could be involved in a quantitative trait is to use the segregating variance in a controlled cross. Assuming a backcross population established by two parental lines fixed for different alleles at relevant loci, a classical estimator of the number of loci underlying the trait is provided by Castle and Wright (*53-55*):

$$n_{\mathrm{CW}} = \frac{|\Delta\overline{Z}|^2}{4\mathrm{Var}[S]} \tag{S3}$$

where $\Delta\overline{Z}$ is the difference of trait means between parental lines, and $\mathrm{Var}[S]$ is the segregating genetic variance of the trait among backcross progeny. In D(DB) and B(BD) females, $\Delta\overline{Z}$ was calculated between backcross females with unintrogressed ($f_Z = 0$) and fully introgressed ($f_Z = 1$) Z chromosomes:

$$\begin{aligned} \Delta\overline{Z}_{\mathrm{D(DB)}} &= \langle W_{\mathrm{D(DB)}} | f_Z = 1 \rangle - \langle W_{\mathrm{D(DB)}} | f_Z = 0 \rangle \\ \Delta\overline{Z}_{\mathrm{B(BD)}} &= \langle W_{\mathrm{B(BD)}} | f_Z = 1 \rangle - \langle W_{\mathrm{B(BD)}} | f_Z = 0 \rangle \end{aligned} \tag{S4}$$

We also assume that environmental variance was constant and no G$\times$E interactions. The environmental variance in each cross was estimated on females with unrecombined Z chromosomes. Then, $\mathrm{Var}[S]$ was estimated by subtracting the environmental variance from the total trait variance.

Since this estimator assumes no linkage among loci, an improved estimator was derived by Zeng (*32*) to incorporate linkage (assuming phenotypic effects are the same for all factors):

$$n_{\mathrm{CWZ}} = \frac{2\bar{c}\, n_{\mathrm{CW}}}{1 - n_{\mathrm{CW}}(1 - 2\bar{c})} \tag{S5}$$

where $\bar{c}$ is the average pairwise recombination probability among all possible loci on the Z chromosome. For our crossover model, this was evaluated as

$$\bar{c} = \frac{1}{2}\int_0^1 \int_0^1 |x - y|\, \mathrm{d}x\, \mathrm{d}y = \frac{1}{6} \tag{S6}$$

[1]To compute $R^2$ when comparing the polygenic model of ovary dysgenesis with observed results, we did not use a logistic mapper. Instead, we coded "Normal" as 1 and all other phenotypes as 0, then performed regression directly on marker ancestry.

Observed and expected genetic variances of pupal weight under a single-QTL and a linear polygenic model are shown in Table S1. Using observed genetic variance, the Castle-Wright-Zeng estimator becomes negative for both backcross directions, which is directly caused by an overly small segregating genetic variance.

##### 1.10 Bayesian QTL model selection

We used the software BayesQTLBIC-1.0-2 (33) to evaluate the posterior probabilities of alternative QTL models. This algorithm assumes additivity among markers. Although the software is extendable to epistasis between markers, the large number of alternative models for epistasis forbids enumerating across all of them. Also, based on the software manual, the prior for epistasis coefficients is not intuitive to define, while the prior for additive effects is well-defined and was set at 0.5 (uniform prior). This limits our implementation of this software to the analysis of only pupal weight.

In pupal weight analysis, we chose 15 sparsely spaced markers on the Z chromosome. This complexity allows us to loop over many models up to six QTLs. The parameters used in the R code is:

```
    result <- bicreg.qtl(x=genotype,y=phenotype,OR=1000000000,maxCol=41,
nbest=500,nvmax=6,prior=0.5,keep.size=1)
```

#### 2. Supplementary Text

##### 2.1 Mathematical details for crossover analysis

**Derivation of the maximum likelihood estimate of crossover frequencies.** Let $n_0$, $n_1$, and $n_2$ be the numbers of haplotypes having zero, one, or two recombination breakpoints on a given chromosome, respectively. Analogously, let $c_0$, $c_1$, and $c_2$ be the probabilities of having zero, one, or two crossovers on that chromosome. We assume at most two crossovers per chromosome. The likelihood of data given parameters is

$$\mathbb{P}(n_0, n_1, n_2 | c_0, c_1, c_2) = \frac{(n_0 + n_1 + n_2)!}{n_0! n_1! n_2!} p_0^{n_0} p_1^{n_1} p_2^{n_2} \tag{S7}$$

where

$$\begin{aligned} p_0 &= c_0 + c_1/2 + c_2/4 \\ p_1 &= c_1/2 + c_2/2 \\ p_2 &= c_2/2 \end{aligned} \tag{S8}$$

The log-likelihood $L(c_1, c_2, c_3) = \ln \mathbb{P}$ is thus given by (the difference is up to a constant):

$$L \sim n_0 \ln\left(1 - \frac{c_1}{2} - \frac{3c_2}{4}\right) + n_1 \ln\left(\frac{c_1}{2} + \frac{c_2}{2}\right) + n_2 \ln\left(\frac{c_2}{4}\right) \tag{S9}$$

Maximizing this function yields

$$\begin{aligned}
c_0 &= (n_0 - n_1 + n_2)/(n_0 + n_1 + n_2) \\
c_1 &= (2n_1 - 4n_2)/(n_0 + n_1 + n_2) \\
c_2 &= 4n_2/(n_0 + n_1 + n_2)
\end{aligned} \tag{S10}$$

However, the data sometimes contained too many recombined individuals (e.g., due to stochastic sampling error) and $c_0$ became negative. Then we maximized $L$ along the boundary of the region $c_1 + c_2 \leq 1$. Since $c_1 > 0$ (no crossover is very unlikely), we substituted $c_1 = 1 - c_2$ into the above likelihood and maximized it within $0 \leq c_2 \leq 1$. This approach produced the following adjustment to the estimate:

$$\begin{aligned}
c_0^* &= 0 \\
c_1^* &= (n_0 - n_2)/(n_0 + n_2) \\
c_2^* &= 2n_2/(n_0 + n_2)
\end{aligned} \tag{S11}$$

#### 2.2 Mathematical details for analyzing pupal weight on the Z chromosome

**Derivation of $R^2_{n\text{-marker}}/R^2_{\text{Z-ancestry}}$ for different architectures of pupal weight.** First, let's clarify our notations. The introgressed ancestry fraction on a chromosome (in this case, the Z chromosome) is denoted as $f$. Marker ancestry at relative position $l$ ($0 \leq l \leq 1$) is $p_l$, and it takes binary values:

$$\begin{cases} 1, & \text{if introgressed} \\ 0, & \text{if not introgressed} \end{cases} \tag{S12}$$

This setup is sufficient for analyzing the single Z chromosome in backcross females.

**Theorem 1** (Statistics under the crossover model)**.** *Crossover on the Z chromosome can be approximated by randomly selecting a position as the only crossover. With this model, we have the following statistics:*

$$\begin{aligned}
\mathbb{E}[f] &= 1/2 \\
\text{Var}[f] &= 1/6 \\
\mathbb{E}[p_l] &= 1/2 \\
\text{Var}[p_l] &= 1/4 \\
\text{Cov}(f, p_l) &= \frac{1}{8}\Big[1 + 2l(1-l)\Big] \\
\text{Cov}(p_{l_1}, p_{l_2}) &= \frac{1}{4}\Big(1 - |l_1 - l_2|\Big)
\end{aligned} \tag{S13}$$

*Proof.* For these statistics, it is helpful to think about the following probabilities on a backcross female's Z chromosome ($\mathrm{d}f$ is the differential in $f$):

$$
\begin{aligned}
&\mathbb{P}(f=0)=\mathbb{P}(f=1)=\frac{1}{4} \quad \text{(Non-recombined)} \\
&\mathbb{P}(\text{Introgressed from the right-hand-side with a fraction } f)=\frac{1}{4}\,\mathrm{d}f \quad \text{(Recombined)} \\
&\mathbb{P}(\text{Introgressed from the left-hand-side with a fraction } f)=\frac{1}{4}\,\mathrm{d}f \quad \text{(Recombined)}
\end{aligned}
\tag{S14}
$$

We immediately have $\mathbb{E}[f]$, $\mathbb{E}[p_l]$, $\mathrm{Var}[f]$, and $\mathrm{Var}[p_l]$ from such probabilities. For covariances,

$$
\begin{aligned}
\mathbb{E}[p_{l_1}p_{l_2}] &= \frac{1}{4}+\frac{1}{4}(1-|l_1-l_2|) \\
\mathbb{E}[f p_l] &= \frac{1}{4}+\frac{1}{4}\int_l^1 f\,\mathrm{d}f+\frac{1}{4}\int_0^l (1-f)\,\mathrm{d}f=\frac{1}{8}\left[3+2l(1-l)\right]
\end{aligned}
\tag{S15}
$$

These quantities are sufficient to derive the covariances between variables. □

**Definition 1** (The linear polygenic model)**.** *For pupal weight $W$, we define its polygenic model as a linear function of average introgressed ancestry ($f$) on the Z chromosome:*

$$W=\alpha f+w \tag{S16}$$

*where $\alpha$ and $w$ are slope and intercept, respectively.*

**Note:** This linear polygenic model will be generalized to nonlinear functions of $f$ when analyzing ovary dysgenesis (discussed in section 2.3). All results in this section must be understood as the joint consequences of:

- A specific genotype-phenotype map (e.g., a linear polygenic model or other QTL models)
- A particular crossover process occurring on these butterflies' Z chromosome
- A backcross brood

With these assumptions in mind, we can predict expected patterns of marker-phenotype association.

**Theorem 2** (Polygenic model & 1-marker scans)**.** *Conditioning on the polygenic model of pupal weight, we have the following relationship for 1-marker scans:*

$$\left.\frac{R^2_{1-\mathrm{marker}}}{R^2_{\mathrm{Z-ancestry}}}\right|_{\mathrm{Polygenic}}=\frac{3}{8}\Big[1+2l(1-l)\Big]^2 \tag{S17}$$

*Proof.* The polygenic model of pupal weight posits that phenotype $W$ is a linear function of $f$. Thus, the regression power of 1-marker scans $R^2_{\text{1-marker}}$ at position $l$, relative to the power of regression using

Z-ancestry $R^2_{\text{Z-ancestry}}$, is simply the squared correlation coefficient between $f$ and $p_l$:

$$\frac{R^2_{\text{1-marker}}}{R^2_{\text{Z-ancestry}}} = \rho^2_{p_l,f} = \frac{\text{Cov}^2(p_l, f)}{\text{Var}[p_l]\text{Var}[f]} = \frac{3}{8}\left[1 + 2l(1-l)\right]^2 \tag{S18}$$

□

The above equation shows an artifactual QTL at the center of the Z chromosome ($l = 0.5$) that maximizes $R^2_{1-\text{marker}}$.

**Theorem 3** (1-QTL model & 1-marker scans)**.** *Conditioning on a 1-QTL model of pupal weight, where the single QTL is at position x, we have the following relationship for 1-marker scans:*

$$\left.\frac{R^2_{1-\text{marker}}}{R^2_{\text{Z}-\text{ancestry}}}\right|_{1-\text{QTL}} = \frac{8}{3}\left[\frac{1 - |l - x|}{1 + 2x(1-x)}\right]^2 \tag{S19}$$

*Proof.* Since the marker at position $x$ contains all phenotypic information, the regression power using another marker at position $l$ is the squared correlation coefficient between $p_x$ and $p_l$, (i.e., $\rho^2_{p_x,p_l}$). Similarly, the regression power using the Z-ancestry is $\rho^2_{p_x,f}$. Thus,

$$\frac{R^2_{1-\text{marker}}}{R^2_{\text{Z}-\text{ancestry}}} = \frac{\rho^2_{p_x,p_l}}{\rho^2_{p_x,f}} = \frac{\text{Cov}^2(p_x, p_l)}{\rho^2_{p_x,f}\text{Var}[p_x]\text{Var}[p_l]} = \frac{8}{3}\left[\frac{1 - |l - x|}{1 + 2x(1-x)}\right]^2 \tag{S20}$$

□

**Theorem 4** (2-QTL model & 1-marker scans)**.** *Conditioning on a 2-QTL model of pupal weight, where the two QTLs are at positions $x_1$ and $x_2$ ($x_1 < x_2$) with equal additive effects, we have the following relationship for 1-marker scans:*

$$\left.\frac{R^2_{1-\text{marker}}}{R^2_{\text{Z}-\text{ancestry}}}\right|_{2-\text{QTL}} = \frac{2}{3}\left[\frac{2 - |l - x_1| - |l - x_2|}{1 + x_1(1-x_1) + x_2(1-x_2)}\right]^2 \tag{S21}$$

*Proof.* Using the same logic as above theorem, this ratio between the two regression powers is

$$\begin{aligned}
\frac{R^2_{1-\text{marker}}}{R^2_{\text{Z}-\text{ancestry}}} &= \frac{\rho^2_{p_{x_1}+p_{x_2},p_l}}{\rho^2_{p_{x_1}+p_{x_2},f}} \\
&= \frac{[\text{Cov}(p_{x_1},p_l)+\text{Cov}(p_{x_2},p_l)]^2}{\text{Var}[p_l]\text{Var}[p_{x_1}+p_{x_2}]} \times \frac{\text{Var}[f]\text{Var}[p_{x_1}+p_{x_2}]}{[\text{Cov}(p_{x_1},f)+\text{Cov}(p_{x_2},f)]^2} \\
&= \frac{\text{Var}[f]}{\text{Var}[p_l]}\left[\frac{\text{Cov}(p_{x_1},p_l)+\text{Cov}(p_{x_2},p_l)}{\text{Cov}(p_{x_1},f)+\text{Cov}(p_{x_2},f)}\right]^2 \\
&= \frac{2}{3}\left[\frac{2-|l-x_1|-|l-x_2|}{1+x_1(1-x_1)+x_2(1-x_2)}\right]^2
\end{aligned} \tag{S22}$$

□

The above 2-QTL/1-marker relationship shows that markers between $x_1$ and $x_2$ all have the same predictive power, because $2-|l-x_1|-|l-x_2| = 2+x_1-x_2$ is independent of $l$ when $x_1 < l < x_2$.

Below, we derive additional results when more than one markers are used.

**Theorem 5** (Polygenic model & $n$-marker scans)**.** *Conditioning on the polygenic model of pupal weight, and assume that n markers with additive effects are used to fit the genotype-phenotype map, the relationship between regression powers is:*

$$\left.\frac{R^2_{n-\text{marker}}}{R^2_{\text{Z}-\text{ancestry}}}\right|_{\text{Polygenic}} = \begin{bmatrix} \rho_{p_{l_1},f} \\ \rho_{p_{l_2},f} \\ \vdots \\ \rho_{p_{l_n},f} \end{bmatrix}^\top \begin{bmatrix} 1 & \rho_{p_{l_1},p_{l_2}} & \cdots & \rho_{p_{l_1},p_{l_n}} \\ \rho_{p_{l_2},p_{l_1}} & 1 & \cdots & \rho_{p_{l_2},p_{l_n}} \\ \vdots & \vdots & \ddots & \vdots \\ \rho_{p_{l_n},p_{l_1}} & \rho_{p_{l_n},p_{l_2}} & \cdots & 1 \end{bmatrix}^{-1} \begin{bmatrix} \rho_{p_{l_1},f} \\ \rho_{p_{l_2},f} \\ \vdots \\ \rho_{p_{l_n},f} \end{bmatrix} \tag{S23}$$

*Proof.* This relationship is by definition the formula for the coefficient of multiple correlation between $f$ and $p_{l_1},\cdots,p_{l_n}$. □

**Corollary 1** (Polygenic model & 2-marker scans)**.** *This is the explicit formula for Theorem 5 using two additive markers ($n=2$):*

$$\left.\frac{R^2_{2-\text{marker}}}{R^2_{\text{Z}-\text{ancestry}}}\right|_{\text{Polygenic}} = \frac{6|l_1-l_2|(l_1+l_2-1)^2+3[1+2l_1(1-l_1)][1+2l_2(1-l_2)]}{8-4|l_1-l_2|} \tag{S24}$$

Equation S24 shows that the two most informative markers under the polygenic model and 2-marker scans are located near $l_1 \approx 0.27$ and $l_2 \approx 0.73$—about a quarter into the chromosome from both ends.

#### 2.3 Mathematical details for analyzing ovary dysgenesis on the Z chromosome

**Artifactual QTL in 1-marker scans when the architecture is polygenic.** The same notation near Equation S12 is used throughout this subsection. In pupal weight analysis, the polygenic model posits that weight is a linear function of introgressed ancestry fraction on the Z chromosome (i.e., $W$ and $f$ are perfectly linearly correlated, ignoring noise). Since the expected ovary phenotypes in D(DB) females and *Heliconius* females are nonlinear with respect to $f$, we now consider a generalized polygenic model, where the expected phenotype $V$ is a continuous function of $f$:

$$V = g(f) \tag{S25}$$

When $g$ is a linear function, we recover the polygenic model for pupal weight. If $g$ is a nonlinear function, it corresponds to global epistasis on Z-linked introgression.

The regression power of a 1-marker scan using the marker at position $l$ against phenotype $V$ is

$$R^2_{1-\text{marker}} = \frac{\text{Cov}^2(p_l, V)}{\text{Var}[p_l]\text{Var}[V]} \tag{S26}$$

Since $\text{Var}[p_l] = 1/4$ and $\text{Var}[V]$ is independent of $l$, the magnitude of $R^2_{1-\text{marker}}$ on different markers depends only on $\text{Cov}(p_l, V)$.

**Theorem 6** (Covariance between $p_l$ and $V$)**.** *Let $h(f) = g(1-f) - g(f)$. The covariance between $p_l$ and $V$ is given by the following formula:*

$$\text{Cov}(p_l, V) = \frac{1}{8}\left[h(0) + \int_0^{1-l} h(f)\,\mathrm{d}f + \int_0^{l} h(f)\,\mathrm{d}f\right] \tag{S27}$$

*Proof.* First, we have

$$\mathbb{E}[p_l] = \frac{1}{2}, \quad \mathbb{E}[V] = \frac{1}{4}g(0) + \frac{1}{4}g(1) + \frac{1}{2}\int_0^1 g(f)\,\mathrm{d}f \tag{S28}$$

The expectation of the product variable $p_l V$ is

$$\mathbb{E}[p_l V] = \frac{1}{4}g(1) + \frac{1}{4}\int_l^1 g(f)\,\mathrm{d}f + \frac{1}{4}\int_{1-l}^1 g(f)\,\mathrm{d}f \tag{S29}$$

Thus,

$$\text{Cov}(p_l, V) = \frac{1}{8}\left[g(1) - g(0) + 2\int_l^1 g(f)\,\mathrm{d}f + 2\int_{1-l}^1 g(f)\,\mathrm{d}f - 2\int_0^1 g(f)\,\mathrm{d}f\right] \tag{S30}$$

Note that the last integral in the bracket can be re-written as follows:

$$2\int_0^1 g(f)\,\mathrm{d}f = \int_0^l g(f)\,\mathrm{d}f + \int_l^1 g(f)\,\mathrm{d}f + \int_0^{1-l} g(f)\,\mathrm{d}f + \int_{1-l}^1 g(f)\,\mathrm{d}f \tag{S31}$$

Substituting the above identity into the covariance formula gives the following result:

$$\mathrm{Cov}(p_l, V) = \frac{1}{8}\left\{g(1) - g(0) + \int_0^{1-l} [g(1-f) - g(f)]\,\mathrm{d}f + \int_0^{l} [g(1-f) - g(f)]\,\mathrm{d}f\right\} \tag{S32}$$

It is thus natural to define $h(f) = g(1-f) - g(f)$, which measures the level of asymmetry of the function $g(f)$ with respect to $f = 1/2$. □

**Theorem 7** (The existence of artifactual QTL in 1-marker scans)**.** *Suppose the polygenic model is true, and the crossover process is as inferred. In that case, the necessary and sufficient condition for a non-zero association between a marker and a trait in a backcross brood is that $g(f)$ is a reflectionally asymmetric function with respect to $f = 1/2$. (Example: Figure S12A-F)*

**Note.** For simplicity, "with respect to" is written as "w.r.t."

*Proof.* The equivalent statement of the theorem is:

$$\text{i) } \mathrm{Cov}(p_l, V) = 0 \text{ for all } l \quad \Leftrightarrow \quad \text{ii) } g(f) \text{ is symmetric w.r.t. } f = 1/2.$$

Proving ii) ⇒ i) is straightforward because a symmetric $g(f)$ means $h(f) \equiv 0$, so $\mathrm{Cov}(p_l, V) \equiv 0$.

To prove i) ⇒ ii), since $\mathrm{Cov}(p_l, V) \equiv 0$, we have

$$0 \equiv \partial_l \mathrm{Cov}(p_l, V) = \frac{1}{8}\{-[g(l) - g(1-l)] + [g(1-l) - g(l)\} \tag{S33}$$

Thus, $g(l) \equiv g(1-l)$, and $g$ is symmetric w.r.t. position $f = 0.5$.

Finally, take the contrapositive statement to get the original theorem:

$$\text{i) } \mathrm{Cov}(p_l, V) = 0 \text{ for some } l \quad \Leftrightarrow \quad \text{ii) } g(f) \text{ is asymmetric w.r.t. } f = 1/2.$$

□

The expected regression power $R^2_{1-\text{marker}}$ will always be a symmetric function w.r.t. $l = 0.5$, because $\mathrm{Cov}(p_l, V) \equiv \mathrm{Cov}(p_{1-l}, V)$. Thus, all properties of $R^2_{1-\text{marker}}$ can be discussed assuming that $l \leq 1/2$. Next, we give a sufficient condition for the existence of a unique peak of $R^2_{1-\text{marker}}$ at the chromosome center.

**Theorem 8** (A sufficient condition for a unique peak of $R^2_{1-\text{marker}}$ at the chromosome center)**.** *If $h(f)$ is a continuous function and has no zeros in $0 < f < 1/2$, then there is a unique peak for $R^2_{1-\text{marker}}$ at the chromosome center ($l = 1/2$). (Examples: Figure S12C,D)*

*Proof.* Again, note that $R^2_{1-\text{marker}}$ is proportional to $\mathrm{Cov}^2(p_l, V)$ by a constant factor, so we only need to prove the existence of a unique peak for $\mathrm{Cov}^2(p_l, V)$ at $l = 1/2$. Second, $h(f)$ is anti-symmetric w.r.t.

$f = 1/2$. If $r < 1/2$, the first integral in Equation S27 is:

$$\int_0^{1-l} h(f)\,\mathrm{d}f = \int_0^{l} h(f)\,\mathrm{d}f \tag{S34}$$

Thus,

$$\mathrm{Cov}^2(p_l, V) = \frac{1}{64}\left[h(0) + 2\int_0^{l} h(f)\,\mathrm{d}f\right]^2 \tag{S35}$$

By anti-symmetry, $h(1/2) = 0$. Since $h(f)$ is continuous and has no zeros in $0 < f < 1/2$, $h(f)$ does not switch sign in $0 < f < 1/2$. Thus, if $h(0) > 0$, the integrand in the previous equation will be positive, and $\left[h(0) + 2\int_0^{l} h(f)\,\mathrm{d}f\right]^2$ is an increasing function of $l$ up to $l = 1/2$. If $h(0) < 0$, the integrand will be negative, and $\left[h(0) + 2\int_0^{l} h(f)\,\mathrm{d}f\right]^2$ is still an increasing function of $l$. If $h(0) = 0$, $h(f)$ will always be positive or negative, and the same result holds. Thus, $\mathrm{Cov}^2(p_l, V)$ is an increasing function of $l$ when $0 \le l \le 1/2$, and by symmetry of $\mathrm{Cov}(p_l, V)$, $\mathrm{Cov}^2(p_l, V)$ has a unique maximum at $l = 1/2$. □

For ovary dysgenesis in D(DB) females and *Heliconius*, since more normal phenotypes are suppressed in backcrosses when the Z chromosome is not recombined in ancestry, we may assume that $h(0) = g(1) - g(0) = 0$. Then,

$$\begin{aligned}
\mathrm{Cov}^2(p_l, V) &= \frac{1}{16}\left[\int_0^{l} h(f)\,\mathrm{d}f\right]^2 \\
\mathrm{Var}[p_l] &= \frac{1}{4} \\
\mathrm{Var}[V] &= \frac{1}{2}\int_0^{1} [g(f) - g(0)]^2\,\mathrm{d}f - \frac{1}{4}\left[\int_0^{1} [g(f) - g(0)]\,\mathrm{d}f\right]^2
\end{aligned} \tag{S36}$$

Without loss of generality, define $\widetilde{g}(f) = g(f) - g(0)$, and so $\widetilde{h}(f) = h(f)$. The regression power can thus be expressed as

$$R^2_{1-\mathrm{marker}}(l) = \left[\int_0^{l} \widetilde{h}(f)\,\mathrm{d}f\right]^2 \Big/ \left\{2\int_0^{1} \widetilde{g}^2(f)\,\mathrm{d}f - \left[\int_0^{1} \widetilde{g}(f)\,\mathrm{d}f\right]^2\right\} \tag{S37}$$

**Artifactual epistatic QTL in 2-marker scans when the architecture is polygenic.** To investigate the statistical interaction between a pair of markers to predict a trait, it is easier to work with binary ancestry defined as

$$\begin{cases} 1, & \text{if introgressed} \\ -1, & \text{if not introgressed} \end{cases} \tag{S38}$$

Note that this representation does not change any prior results assuming additivity among markers. For two markers at positions $l_1$ and $l_2$, we assume that $l_1 \le l_2$. The following statistics are associated with the crossover model with the new notation:

$$\begin{aligned}\mathbb{E}[p_{l_1}p_{l_2}] &= 1-(l_2-l_1)\\ \mathrm{Var}[p_{l_1}p_{l_2}] &= (l_2-l_1)(2-l_2+l_1)\end{aligned} \tag{S39}$$

Let the average magnitude of $g(f)$ be:

$$\overline{g} = \int_0^1 g(f)\,\mathrm{d}f \tag{S40}$$

Then, we have the covariance between $p_{l_1}p_{l_2}$ and $V$ as follows.

**Theorem 9** (Covariance between $p_{l_1}p_{l_2}$ and $V$). *Let $H(f) = g(f) + g(1-f) - 2\overline{g}$. The covariance between $p_{l_1}p_{l_2}$ and $V$ is given by the following formula:*

$$\mathrm{Cov}(p_{l_1}p_{l_2}, V) = \int_{l_1}^{l_2} \left[\frac{1}{4}H(0) - \frac{1}{2}H(f)\right] \mathrm{d}f \tag{S41}$$

*Proof.* First, we arrange terms into the form of $g(f)+g(1-f)$:

$$\begin{aligned}\mathbb{E}[p_{l_1}p_{l_2}V] &= \frac{1}{4}g(1) + \frac{1}{4}g(0) + \frac{1}{4}\left(\int_0^{l_1} + \int_{l_2}^{1} - \int_{l_1}^{l_2}\right) g(f)\,\mathrm{d}f + \frac{1}{4}\left(\int_0^{1-l_2} + \int_{1-l_1}^{1} - \int_{1-l_2}^{1-l_1}\right) g(f)\,\mathrm{d}f\\ &= \frac{1}{4}g(1) + \frac{1}{4}g(0) + \frac{1}{4}\left(\int_0^{l_1} + \int_{l_2}^{1} - \int_{l_1}^{l_2}\right) g(f)\,\mathrm{d}f + \frac{1}{4}\left(\int_{l_2}^{1} + \int_0^{l_1} - \int_{l_1}^{l_2}\right) g(1-f)\,\mathrm{d}f\\ &= \frac{1}{4}\left[g(1) + g(0) + \left(\int_0^{l_1} + \int_{l_2}^{1} - \int_{l_1}^{l_2}\right)[g(f)+g(1-f)]\,\mathrm{d}f\right]\\ &= \frac{1}{4}\left[H(0) + \left(\int_0^{l_1} + \int_{l_2}^{1} - \int_{l_1}^{l_2}\right)H(f)\,\mathrm{d}f\right] + (1+l_1-l_2)\overline{g}\\ \mathbb{E}[p_{l_1}p_{l_2}]\mathbb{E}[V] &= (1+l_1-l_2)\left[\frac{1}{4}H(0) + \overline{g}\right]\end{aligned} \tag{S42}$$

Thus,

$$\begin{aligned}\mathrm{Cov}(p_{l_1}p_{l_2}, V) &= \mathbb{E}[p_{l_1}p_{l_2}V] - \mathbb{E}[p_{l_1}p_{l_2}]\mathbb{E}[V]\\ &= \frac{1}{4}\left(\int_0^{l_1} + \int_{l_2}^{1} - \int_{l_1}^{l_2}\right)H(f)\,\mathrm{d}f + \frac{1}{4}(l_2-l_1)H(0)\\ &= \frac{1}{4}\left(\int_0^{1} - 2\int_{l_1}^{l_2}\right)H(f)\,\mathrm{d}f + \frac{1}{4}\int_{l_1}^{l_2} H(0)\,\mathrm{d}f\\ &= \frac{1}{4}\int_0^1 H(f)\,\mathrm{d}f + \int_{l_1}^{l_2}\left[\frac{1}{4}H(0) - \frac{1}{2}H(f)\right]\mathrm{d}f\end{aligned} \tag{S43}$$

Since $\int_0^1 H(f)\,\mathrm{d}f = 0$, we have

$$\mathrm{Cov}(p_{l_1}p_{l_2}, V) = \int_{l_1}^{l_2} \left[\frac{1}{4}H(0) - \frac{1}{2}H(f)\right] \mathrm{d}f \tag{S44}$$

□

**Theorem 10** (The existence of artifactual interacting QTL pairs in 2-marker scans)**.** *Suppose the polygenic model is true, and the crossover process is as inferred. In that case, the necessary and sufficient condition for a non-zero interaction between a pair of markers is that $g(f)$ is a rotationally asymmetric function with respect to the point $(1/2, f(1/2))$ by a degree of 180°. Examples: Figure S14.*

*Proof.* The logic is similar to the proof of Theorem 7. Take the derivative w.r.t. either $l_1$ or $l_2$ of the above covariance, we get:

$$0 \equiv \partial_{l_2}\mathrm{Cov}[p_{l_1}p_{l_2}, V] = \frac{1}{4}H(0) - \frac{1}{2}H(l_2) \tag{S45}$$

Thus, $H(f)$ is a constant function, implying that $g(f) + g(1-f) \equiv$ Const. This relationship indicates that $g$ is a rotationally symmetric function w.r.t. the point $(1/2, g(1/2))$ and the rotation angle $\pi$. Conversely, if $g$ is rotationally symmetric, the integrand becomes zero, and covariance is globally zero. □

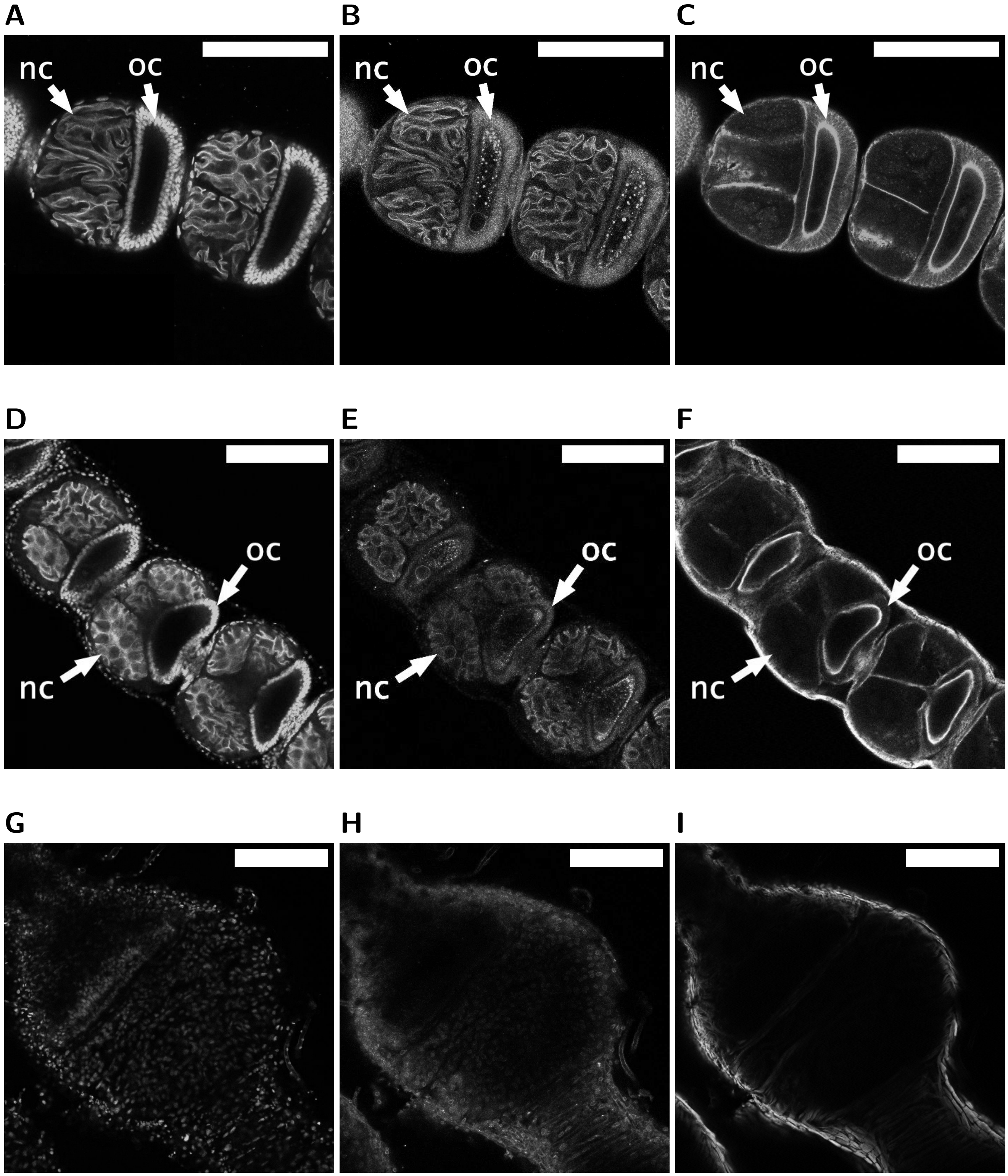

**Figure S1:** Confocal imaging of ovary phenotypes in *Papilio*. Scale bar=200$\mu$m. Left column: Hoechst (DNA); Middle column: WGA (membrane); Right column: Phalloidin (actin filaments). **(A-C)** Phenotype Normal in pure individuals. **(D-F)** Phenotype Normal in $F_1$ DB hybrids. **(G-I)** Phenotype Empty in $F_1$ BD hybrids.

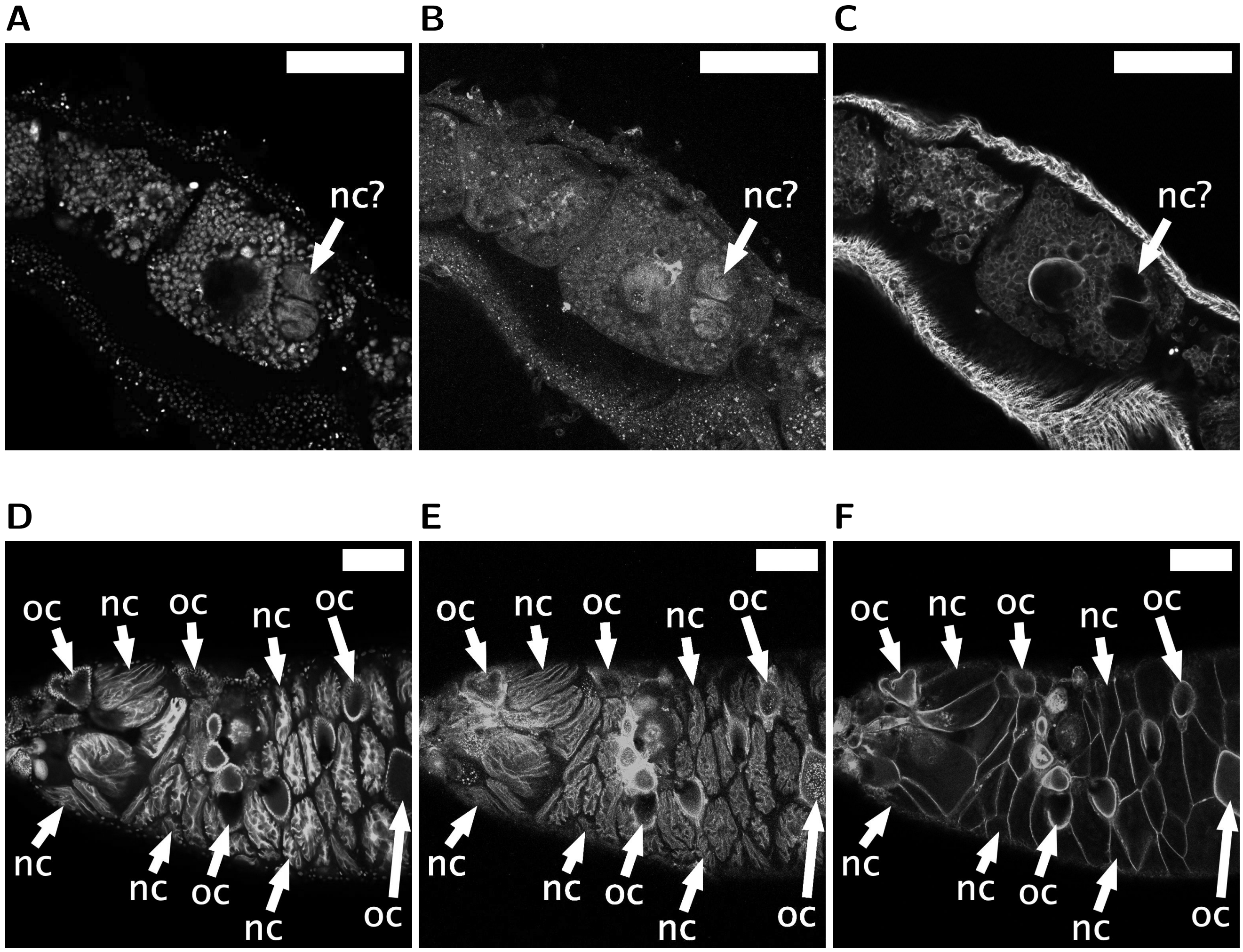

**Figure S2:** Confocal imaging of ovary phenotypes in *Papilio* (continued). Scale bar=200$\mu$m. Left column: Hoechst (DNA); Middle column: WGA (membrane); Right column: Phalloidin (actin filaments). **(A-C)** Phenotype Diminished (only in backcross individuals). **(D-F)** Phenotype Jammed (only in backcross individuals).

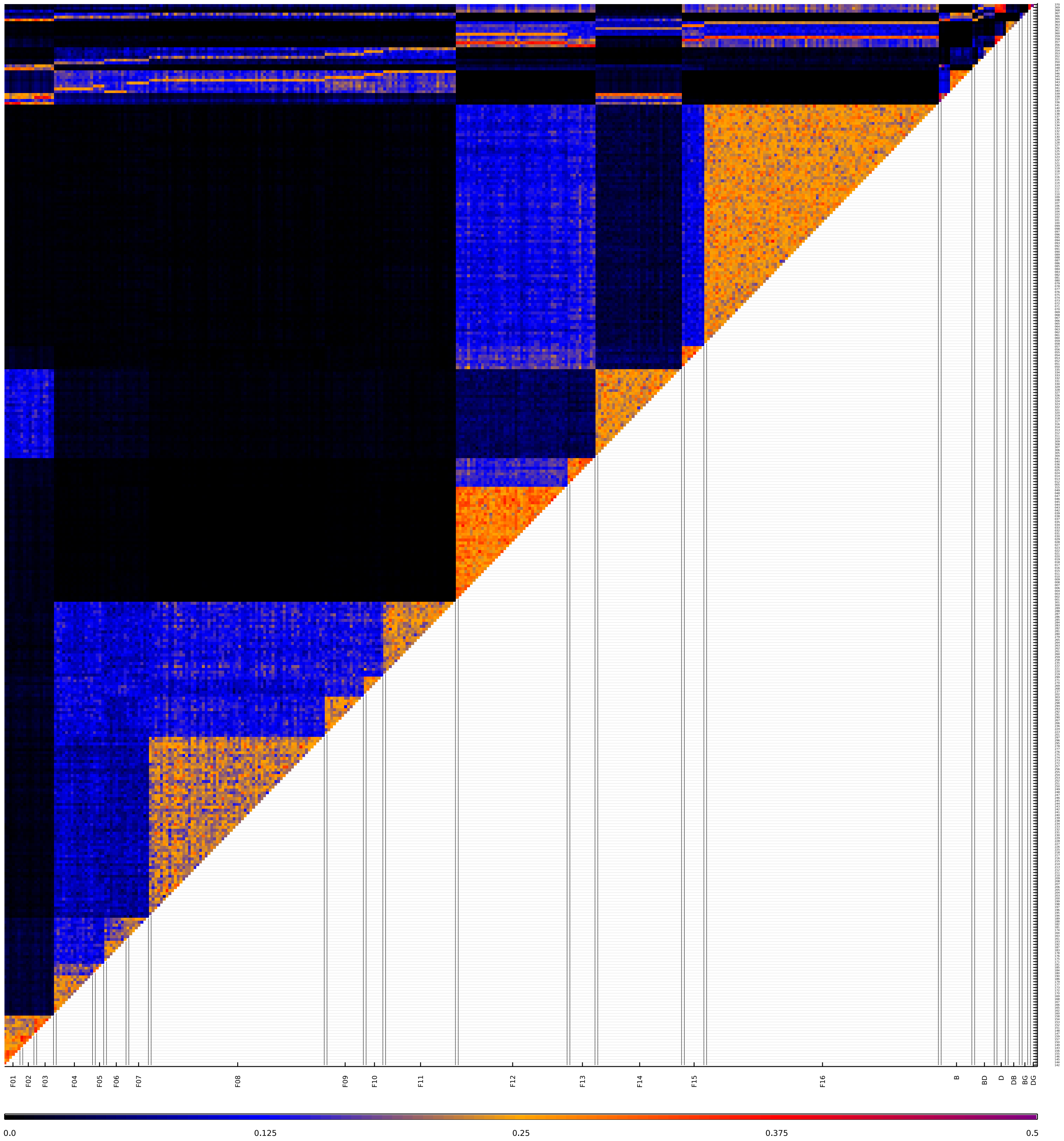

**Figure S3:** Kinship among individuals in *Papilio* after correcting for misplaced individuals in the pedigree. The horizontal axis contains family information (each "FXX" is a single family), and the vertical axis shows individual identifiers.

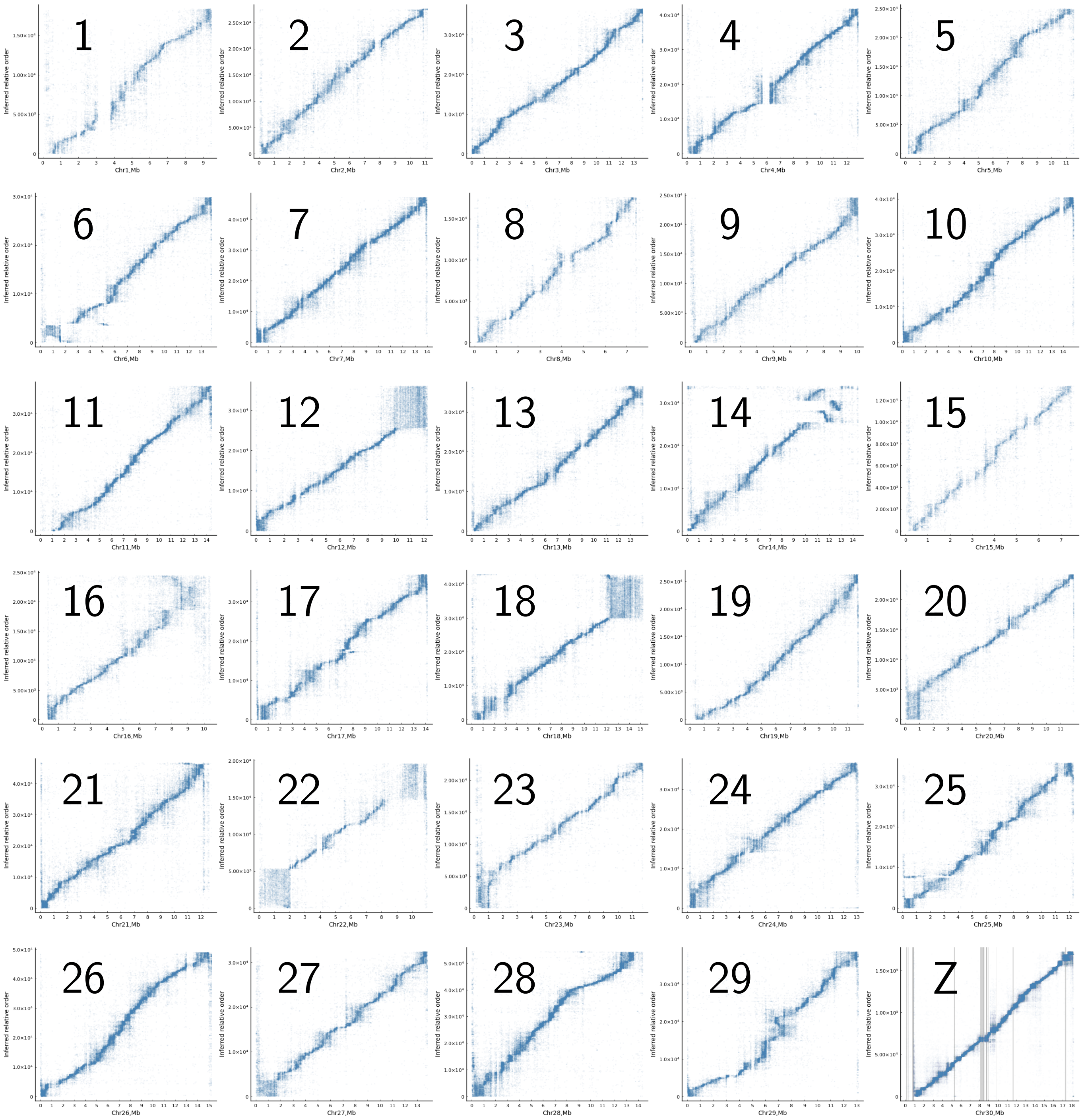

**Figure S4:** Inferred de novo marker order against the corrected reference genome of *Papilio bianor* shows good collinearity except for chromosome 14. Vertical lines in gray represent boundaries between PacBio scaffolds. Some chromosomal ends appear to have recombination suppressed (large blocks of unordered markers).

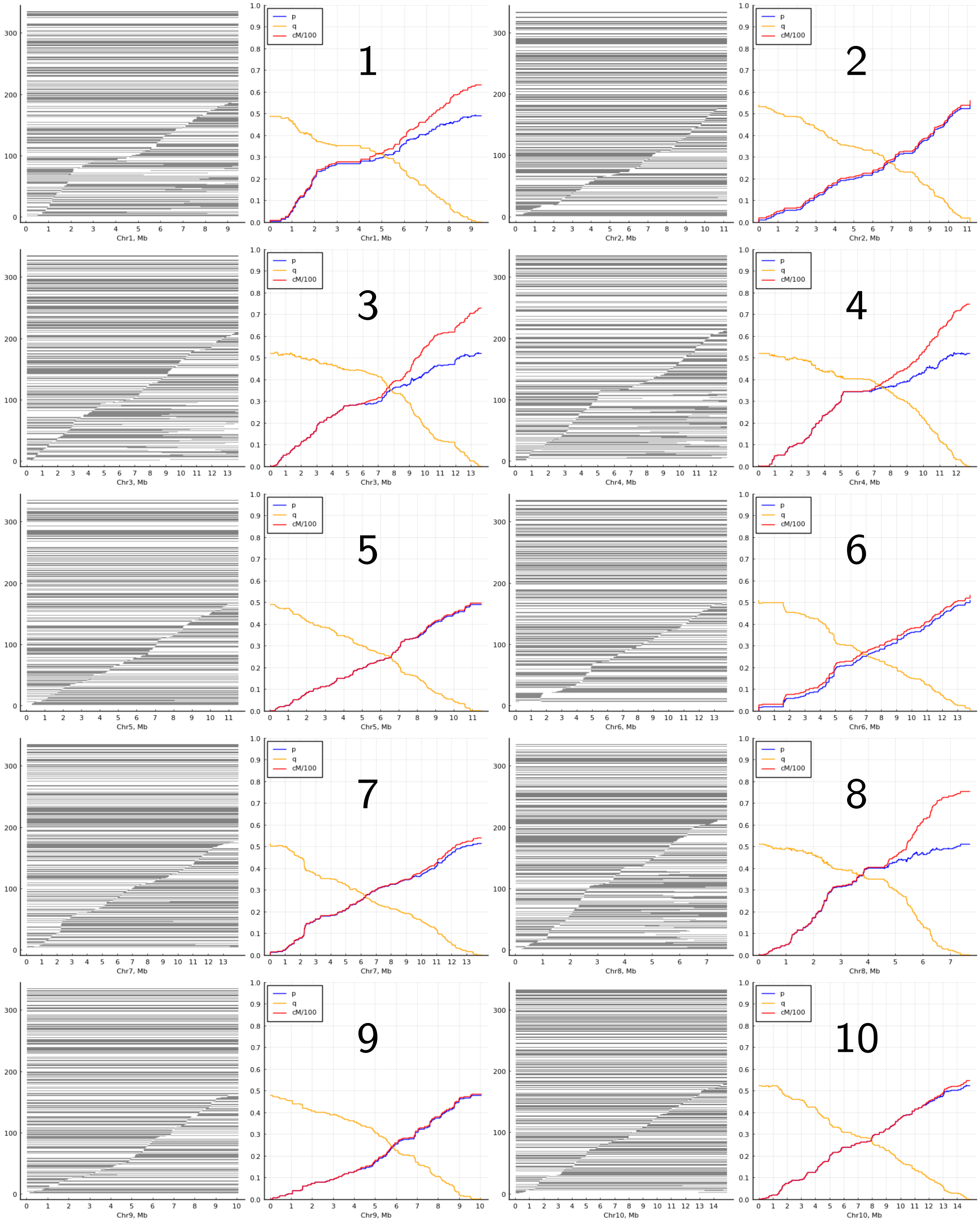

**Figure S5:** Inferred paternal haplotypes among all backcross individuals on chromosomes 1-10 in *Papilio*. Blue curves show the recombination probability of each marker to the left end of each chromosome. Yellow curves show the recombination probability of each marker to the right end of each chromosome. Red curves are linkage maps measured in cM.

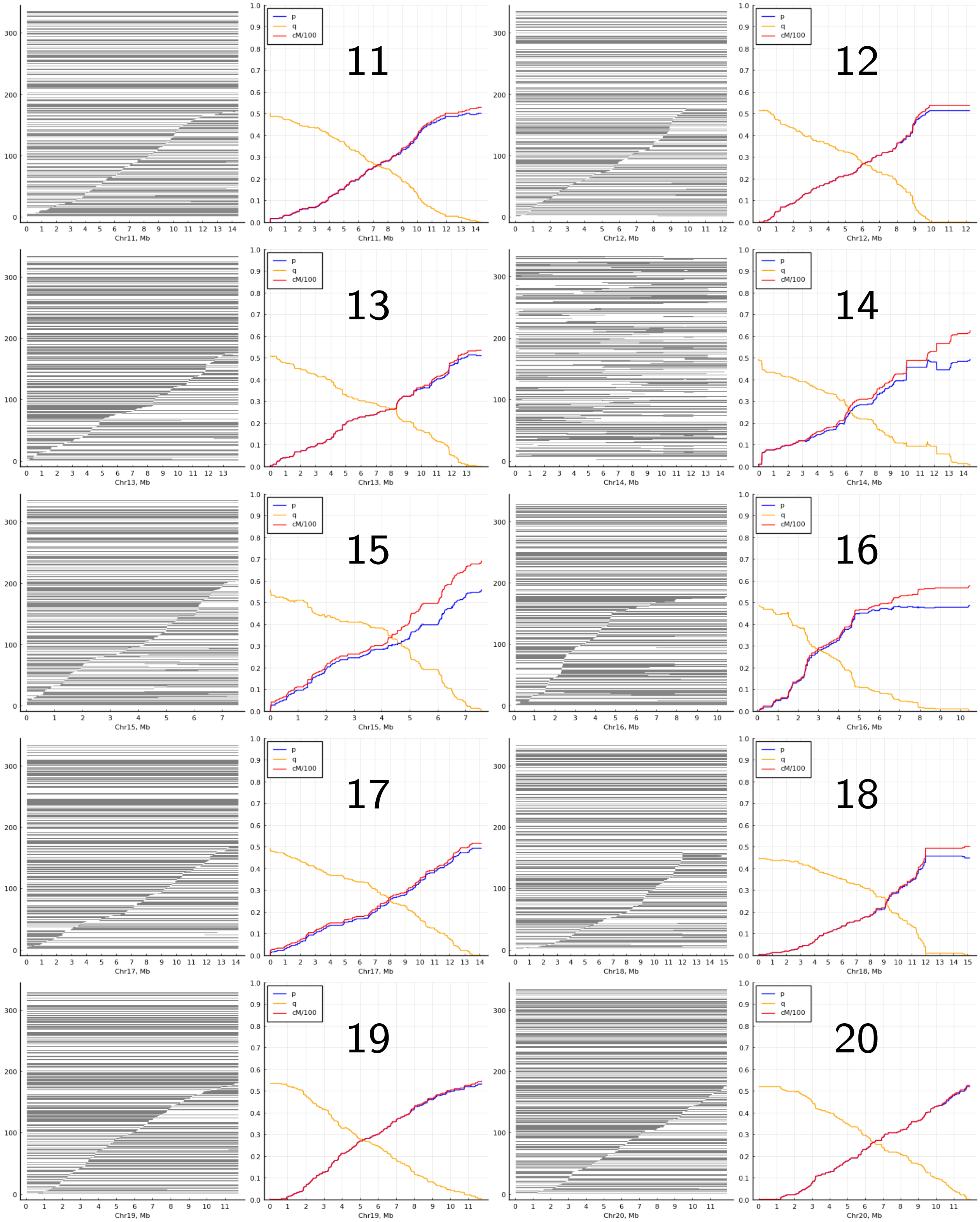

**Figure S6:** Inferred paternal haplotypes among all backcross individuals on chromosomes 11-20 in *Papilio*. Blue curves show the recombination probability of each marker to the left end of each chromosome. Yellow curves show the recombination probability of each marker to the right end of each chromosome. Red curves are linkage maps measured in cM.

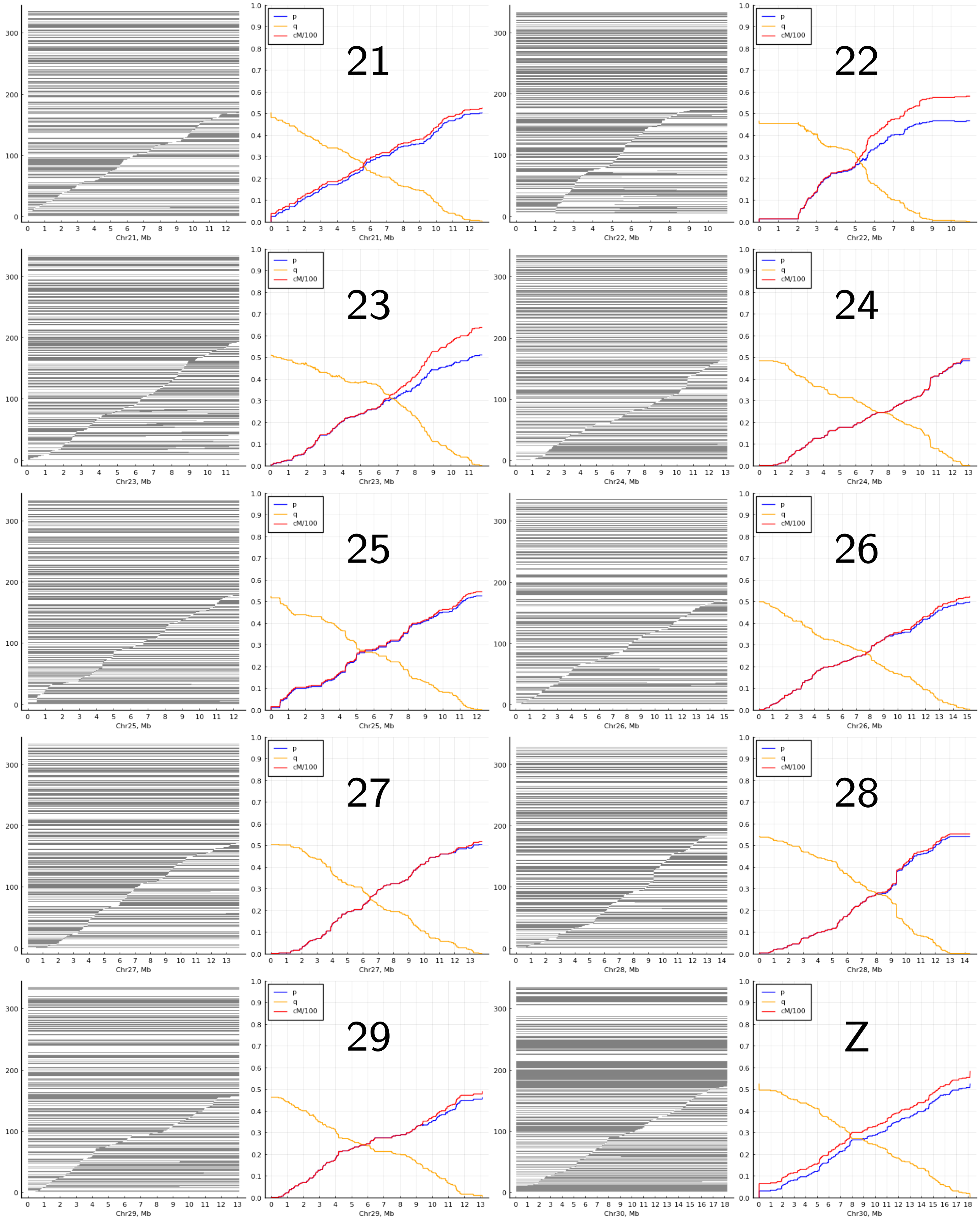

**Figure S7:** Inferred paternal haplotypes among all backcross individuals on chromosomes 21-Z in *Papilio*. Blue curves show the recombination probability of each marker to the left end of each chromosome. Yellow curves show the recombination probability of each marker to the right end of each chromosome. Red curves are linkage maps measured in cM.

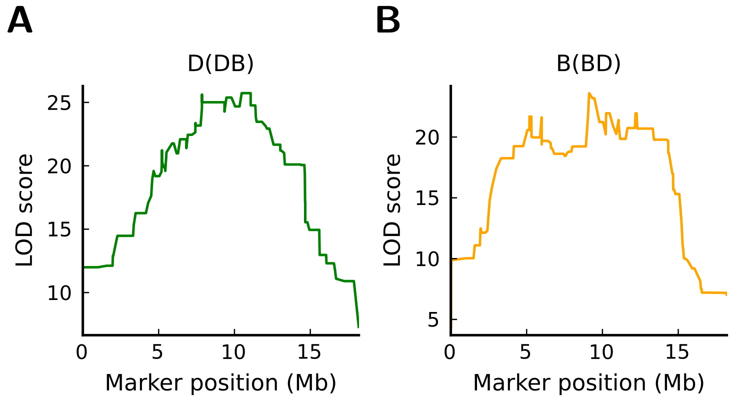

**C**

| Cross | Peak (Mb) | 95% CI (Mb) |
|---|---|---|
| D(DB) | 11.06879 | [7.855832, 11.37611] |
| B(BD) | 9.124007 | [9.124007,12.254113] |

**Figure S8:** One-marker scans of pupal weight in *Papilio* on the Z chromosome by r/qtl2. **(A,B)** LOD scores on the Z chromosome. **(C)** Peaks identified by r/qtl2 and their confidence intervals (CI) at the 95% level.

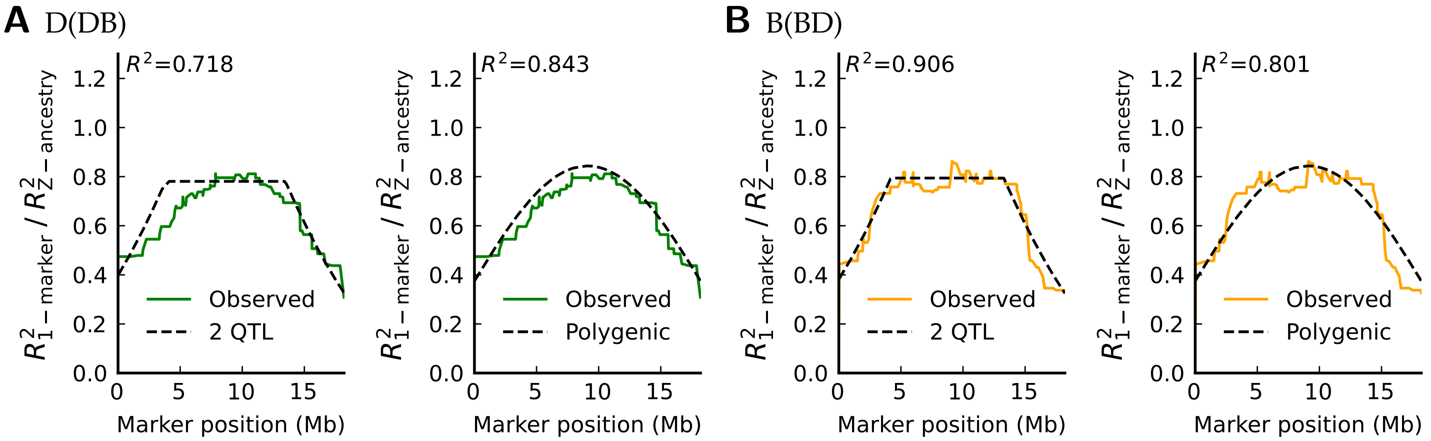

**Figure S9:** Expected results of 1-marker scans with the polygenic model versus 2-QTL models that best fit the observed curves in *Papilio* pupal weight. **(A)** In D(DB) females, the polygenic model can better fit 1-marker scans. For the best 2-QTL model, the relative locations of the two QTLs are 0.21 and 0.74. **(B)** In B(BD) females, the fully polygenic model is worse at fitting 1-marker scans than the best 2-QTL model. For the best 2-QTL model, the relative locations of the two QTLs are 0.23 and 0.73.

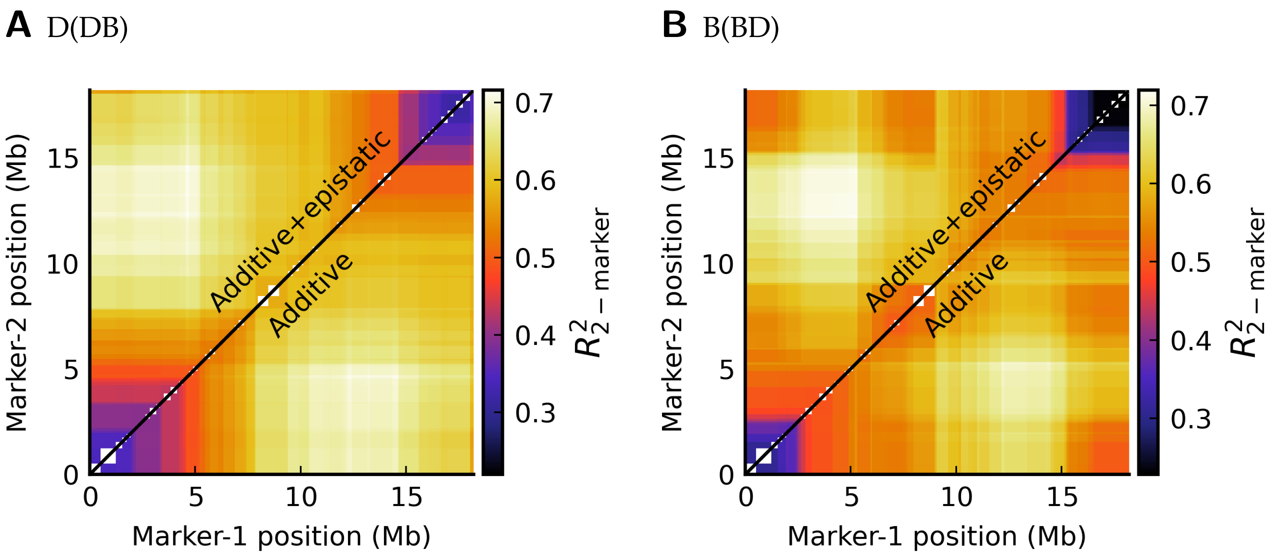

**Figure S10:** Results of 2-marker regression on pupal weight in *Papilio*. Model prediction powers are nearly identical between the additive model ($W \sim p_{l_1} + p_{l_2}$) and the full model with an extra epistasis term ($W \sim p_{l_1} + p_{l_2} + p_{l_1} p_{l_2}$). Thus, epistasis adds little information to predicting pupal weight in backcrosses.

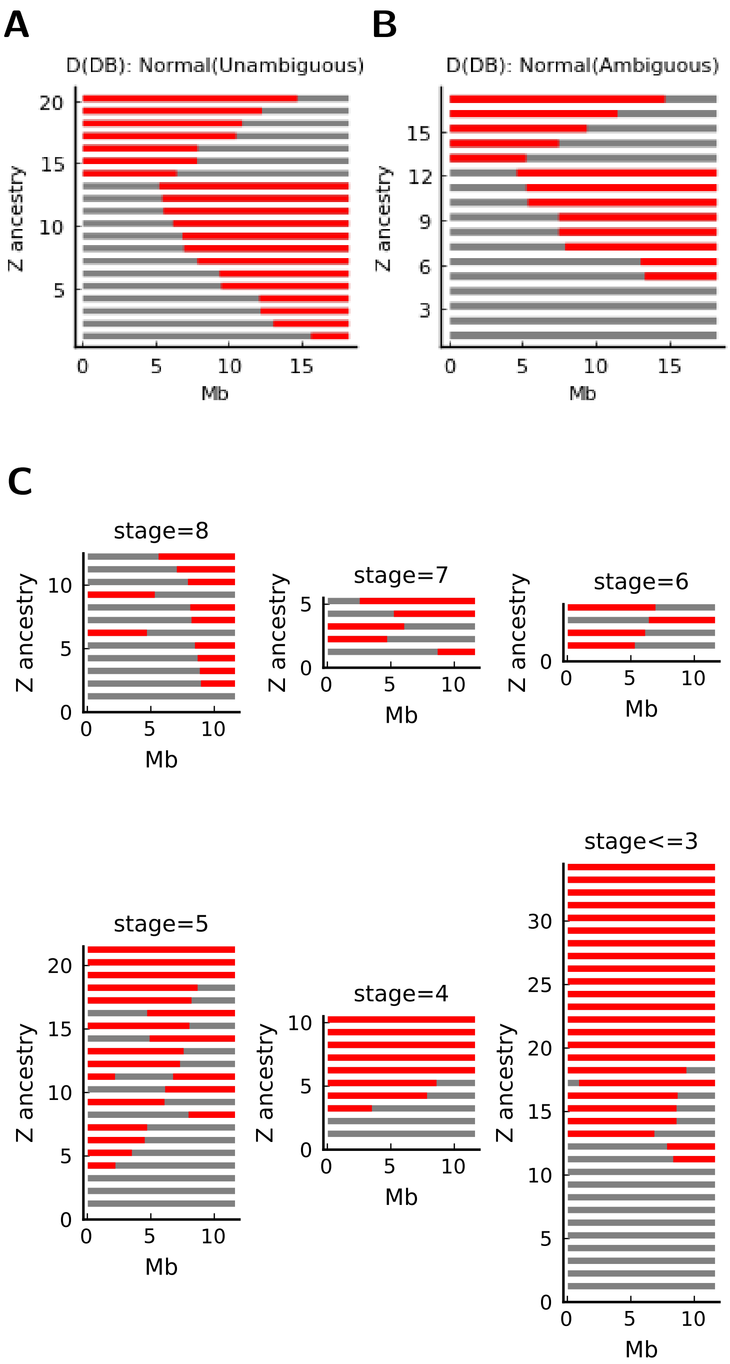

**Figure S11:** Z chromosome ancestry haplotypes in D(DB) females and *Heliconius* backcrosses. **(A,B)** Z chromosome haplotypes in D(DB) females associated with phenotype Normal. This phenotype is associated with Z chromosomes recombined in either direction. Gray: inherited from *P. dehaanii*; Red: inherited from *P. bianor*. **(C)** Z chromosome haplotypes in *Heliconius* backcross females grouped by ovary stages (larger=more normal). Gray: inherited from *H. pardalinus butleri*; Red: inherited from *H. p. sergestus*.

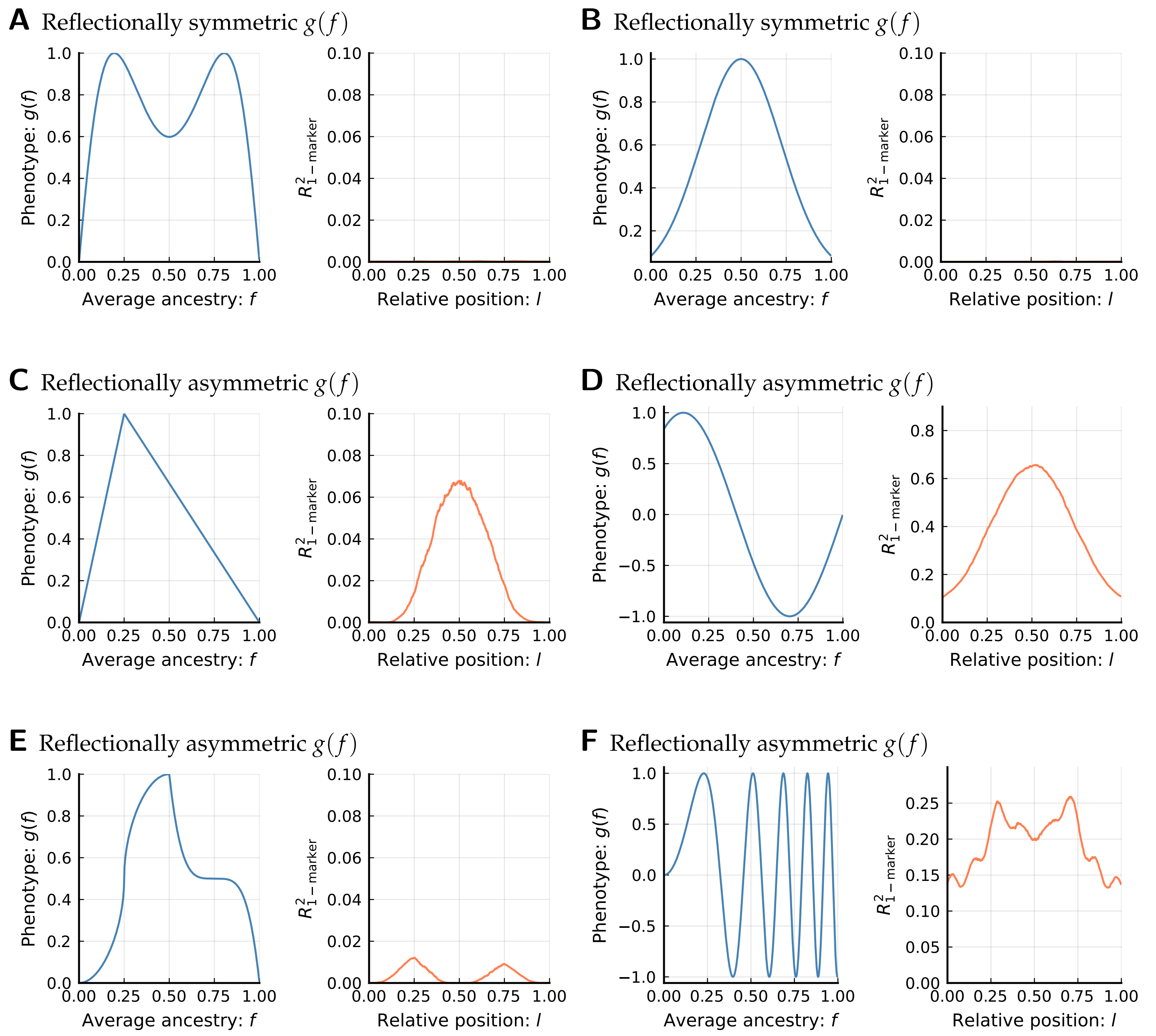

**Figure S12:** Different reflectional symmetry of $g(f)$ leads to different results of 1-marker scans. The uneven distribution of $R^2$ does not reflect an uneven distribution of phenotypic effects, because the model is fully polygenic. Simulated using $10^4$ backcross individuals. **(A,B)** A reflectionally symmetric $g(f)$ w.r.t. $f = 0.5$ produces no marker-phenotype association in 1-marker scans. **(C,D)** A reflectionally asymmetric $g(f)$ satisfying Theorem 8, i.e., no zeros in $h(f)$ when $0 < f < 1/2$, produces a unique peak at the chromosome center. **(E,F)** A reflectionally asymmetric $g(f)$ violating conditions in Theorem 8 can produce multiple peaks in 1-marker scans, but the shape of $R^2$ is still symmetric w.r.t. $l = 1/2$.

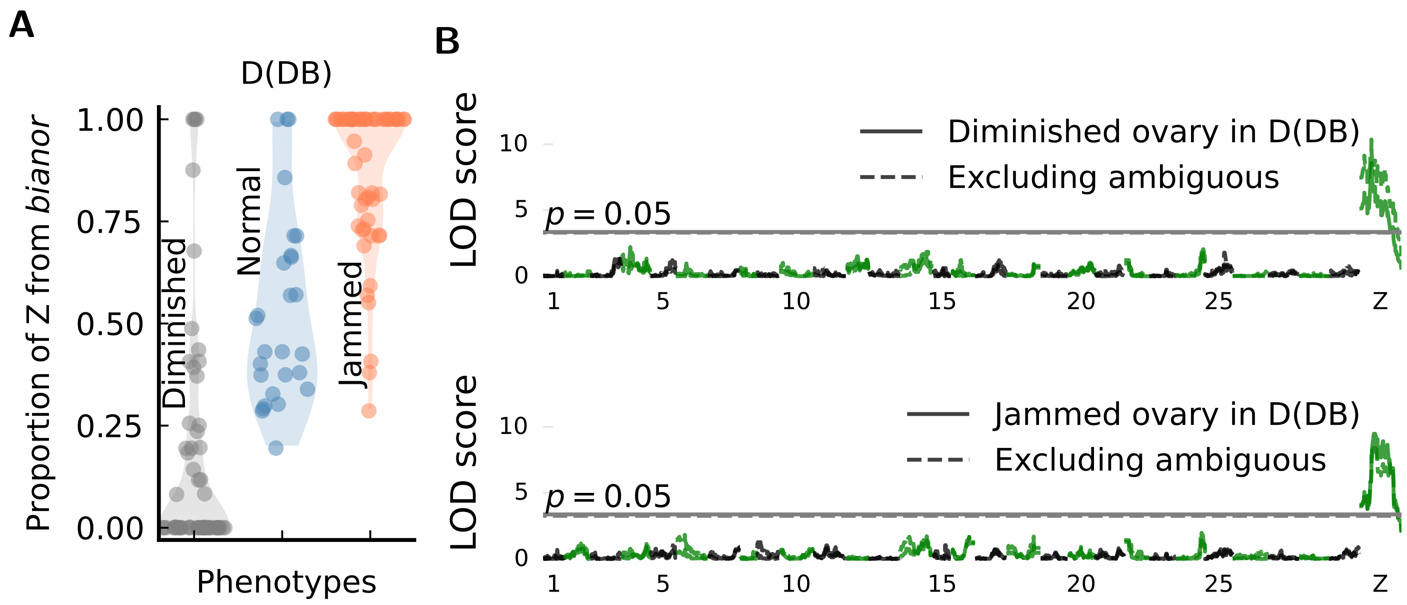

**Figure S13:** Ovary phenotypes in D(DB) females partitioned by the Z-chromosome ancestry fraction. **(A)** Ovary phenotypes in D(DB) females are well described by the Z-chromosome ancestry fraction. Sample dots represent one round of phenotype assignment for individuals with ambiguous phenotypes. **(B)** LOD scores of defective ovary phenotypes in D(DB) females using 1-marker scans. The Z chromosome is significantly associated with phenotypes Diminished and Jammed. This significant association is predicted by the polygenic model, because there is a strong reflectional asymmetry in $g(f)$ when the phenotype occurs only when the Z chromosome has little introgression (Diminished) or with nearly full introgression (Jammed).

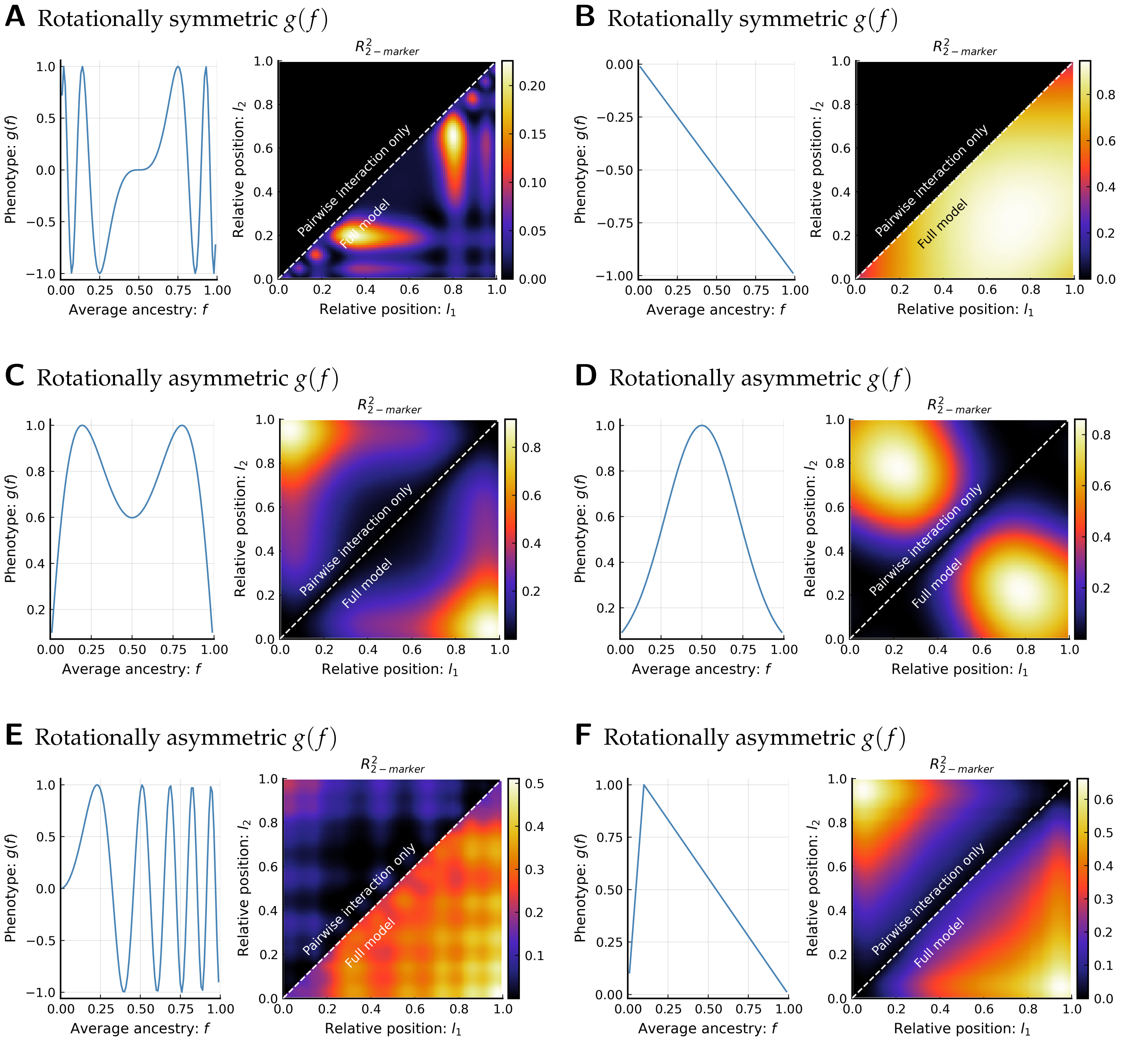

**Figure S14:** Different rotational symmetry of $g(f)$ leads to different results of 2-marker scans. The heatmap contains two genotype-phenotype models. The "Pairwise interaction only" model uses the regression $V \sim 1 + p_{l_1} p_{l_2}$, while the "Full model" includes additive terms: $V \sim 1 + p_{l_1} + p_{l_2} + p_{l_1} p_{l_2}$. The uneven distribution of $R^2$ does not reflect an uneven distribution of phenotypic effects, because the model is fully polygenic. Simulated using $10^4$ backcross individuals. **(A,B)** A rotationally symmetric $g(f)$ w.r.t. the function center produces no interaction between markers. **(C-F)** A rotationally asymmetric $g(f)$ w.r.t. the function center produces interaction between markers. If the function is also reflectionally symmetric (panels C and D), there will be no additive effects, so the interaction term dominates the full model.

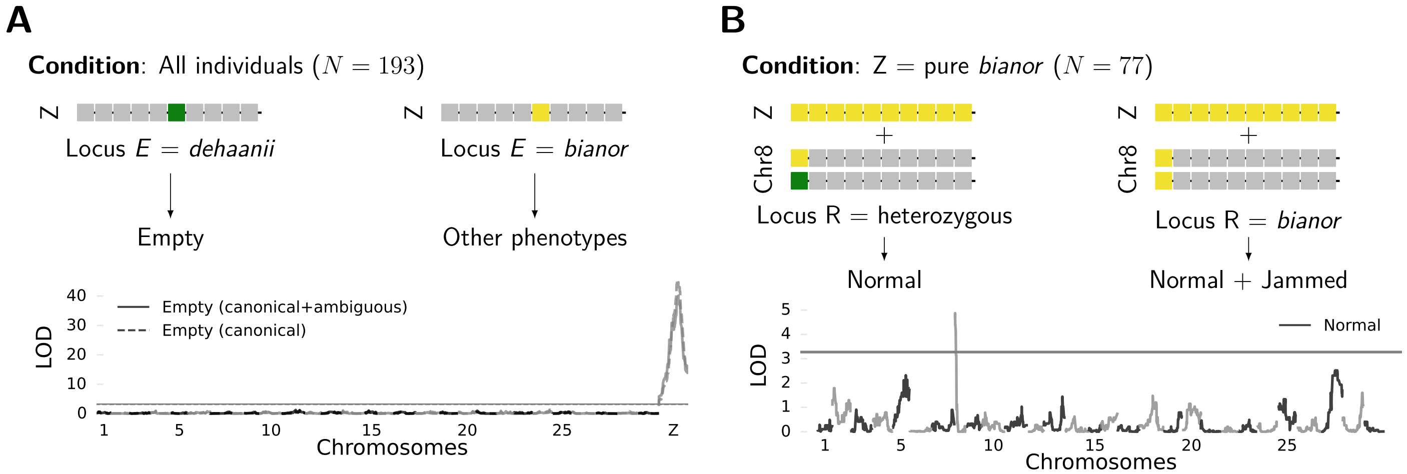

**Figure S15:** Two narrow regions of major effects control ovary dysgenesis in maternally *bianor* hybrids. **(A)** Phenotype Empty is dominantly controlled by Locus $E$ on the Z chromosome. The LOD plot shows both the score for the canonical Empty phenotype as well as the score when we include a few ambiguous individuals that are classified as Empty. **(B)** If the Z chromosome is purely *bianor*, introgression on Locus $R$ from *dehaanii* suppresses abnormal phenotypes.

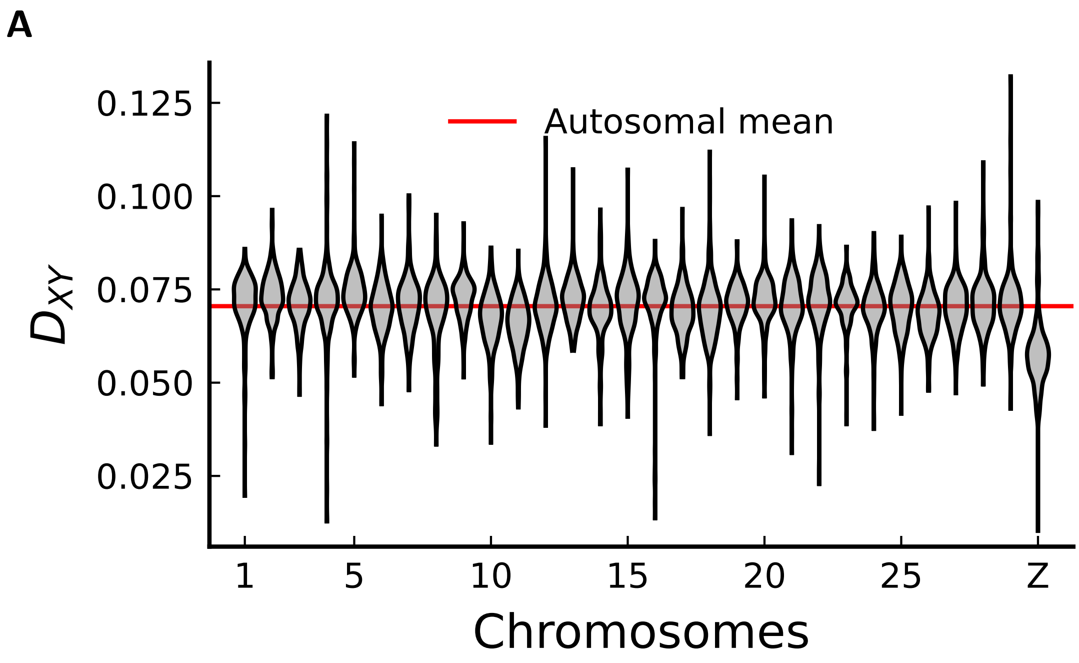

**Figure S16:** Average sequence divergence ($D_{XY}$) between parental *P. dehaanii* and *P. bianor* used in the experiment. Each data point is estimated for 50kb non-overlapping chromosomal windows.

**Table S1:** Genetic variance of pupal weight ($V_g$, unit: gram$^2$) among backcross females in *Papilio* under different architectures. A negative $n_{\mathrm{CWZ}}$ results from a very small genetic variance due to many effective factors.

| Cross direction | D(DB) | B(BD) |
|---|---|---|
| Expected $V_g$ for a single QTL | 0.250 | 0.168 |
| Expected $V_g$ for a linear polygenic model | 0.166 | 0.112 |
| Observed $V_g$ (i.e., $\mathrm{Var}[S]$) | 0.136 | 0.0977 |
| Estimated effective number of loci ($n_{\mathrm{CWZ}}$) | -2.720 | -3.860 |

**Table S2:** Summary of locus E and locus R in *Papilio*

| Locus | Chromosome | Position (Mb) | LOD | 95% Confidence interval (Mb) |
|---|---|---|---|---|
| E (canonical+ambiguous) | Z | 12.18279 | 40.12502 | [11.64904, 12.25411] |
| E (canonical) | Z | 11.45892 | 45.56313 | [11.37611 12.25411] |
| R | 8 | 0.366353 | 4.875207 | [0.005916 0.784124] |

**Table S3:** The ratio of genetic variance between male and female pupal weight among backcross individuals in *Papilio*

| Cross direction | D(DB) | B(BD) |
|---|---|---|
| $V_{g,\mathrm{Male}}/V_{g,\mathrm{Female}}$ | 0.37 | 0.23 |
| 95% Confidence interval of $V_g$ ratio | (0.18, 0.62) | (0.05, 0.46) |
